# ^13^C ENDOR Spectroscopy-Guided MD Computations Reveals the Structure of the Enzyme-Substrate Complex of an Active, N-linked Glycosylated Lipoxygenase

**DOI:** 10.1101/2022.12.07.519351

**Authors:** Ajay Sharma, Chris Whittington, Mohammed Jabed, S. Gage Hill, Anastasiia Kostenko, Tao Yu, Pengfei Li, Brian M. Hoffman, Adam R. Offenbacher

## Abstract

Lipoxygenases (LOXs) are enzymes responsible for producing important cell signaling mediators and have been extensively studied for their potential clinical relevance as well as to advance our understanding of enzyme catalysis. The common inability to capture and characterize LOX-substrate complexes by Xray co-crystallography requires the development of alternative structural methods. We previously reported the integration of ^13^C/^1^H electron nuclear double resonance (ENDOR) spectroscopy and molecular dynamics (MD) to visualize the complex structure of the paradigmatic LOX from soybean, SLO, with substrate linoleic acid (LA). However, this required substitution of the catalytic mononuclear, nonheme iron by the structurally faithful, yet inactive Mn^2+^ ion as a spin-probe. Unlike canonical Fe-LOXs from plants and animals, LOXs from pathogenic fungi contain active mononuclear manganese metallocentres. Here, we report the ground-state active-site structure of the native, fully glycosylated fungal LOX from *M. oryzae, Mo*LOX complexed with LA obtained through the ^13^C/^1^H ENDOR-guided MD approach. The Mn-oxygen-to-LA donor carbon distance (DAD) for *Mo*LOX-LA, 3.4 ± 0.3 Å, matches the distance in the single representative X-ray co-structure of an animal 8R-LOX with its natural fatty acid substrate, and slightly elongated from that of the SLO-LA complex, 3.1 ± 0.2 Å, despite its ‘carboxylate-out’ substrate binding orientation versus ‘carboxylate-in’ for SLO. The results provide unique insight into the evolutionary divergence of the ground-state DAD in the LOX family, which influences the activation barrier for hydrogen tunneling, and give a structural basis for guiding development of *Mo*LOX inhibitors. The work highlights the robustness of ENDOR-guided MD approach to describe LOX-substrate structures that elude conventional X-ray techniques.

## Introduction

Lipoxygenases (LOXs) are a family of enzymes widely represented in plants, animals, fungi and select prokaryotes.^*1-3*^ LOXs oxidize polyunsaturated fatty acids to form a diverse array of potent bioactive cell-signaling mediators. For example,^*1*^ the six human LOX isozymes produce a variety of pro-inflammatory oxylipins linked to chronic inflammation, including atherosclerosis, diabetes, stroke, asthma and select cancers.^*4*^ Despite the clinical importance of these human LOXs, there is only one FDA approved anti-inflammatory drug, Zileuton, that targets human 5-LOX activity linked to asthma.^*5*^, ^*6*^ The dearth of LOX therapeutics is due in part to the lack of high-resolution structural information on the enzyme-substrate (ES) complexes, with only a few representative co-crystal structures available of LOXs with natural or mimetic substrates, or isoform-selective inhibitors.^*7-10*^ This contrasts with cyclooxygenase (COX) enzymes, where the availability of high-resolution structures of protein-inhibitor complexes for both isoforms have aided the development and evolution of non-steroidal anti-inflammatory drugs (NSAIDs).^*6*^

In addition to its medical relevance, the initial C-H bond cleavage associated with the LOX reaction occurs by a non-classical tunneling process, and extensive experimental and theoretical studies have advanced our understanding of the critical link between protein structure, dynamics, and the origins of enzyme catalysis.^*11*^ Thus, implementation of alternative structural methods, complementary to traditional x-ray structural determination, is needed for acquiring detailed information about active protein configurations that can provide a framework to facilitate the design of structure-based inhibitors for therapeutic intervention and to shed additional light onto the structural regulation of enzymatic hydrogen tunneling.

LOXs are also important in pathogenic organisms. Of the ten most pathogenic plant fungi,^*12*^ the rice blast fungus, *M. oryzae*, is considered the most destructive: *M. oryzae* is responsible for the loss of nearly 1/3 of the world’s rice crops, causing the loss of enough food to feed up to 60 million people. The lipoxygenase from *M. oryzae* (*Mo*LOX), the focus of this report, is synthesized at the onset of pathogenesis and is secreted by the fungus along with lipases.^*3*^ *In vitro* studies have shown that *Mo*LOX oxidizes both polyunsaturated fatty acids and phospholipids, generating unique bis-allylic hydroperoxides (**Scheme 1**) that are thought to induce oxidative damage and necrosis of rice leaves.^*3*^, ^*13*^ Thus, it has been proposed as one of the prime agents causing the pathology of rice blast disease.^4^ Development of inhibitors for *Mo*LOX could potentially provide a successful intervention for rice blast disease.

**Scheme 1.**
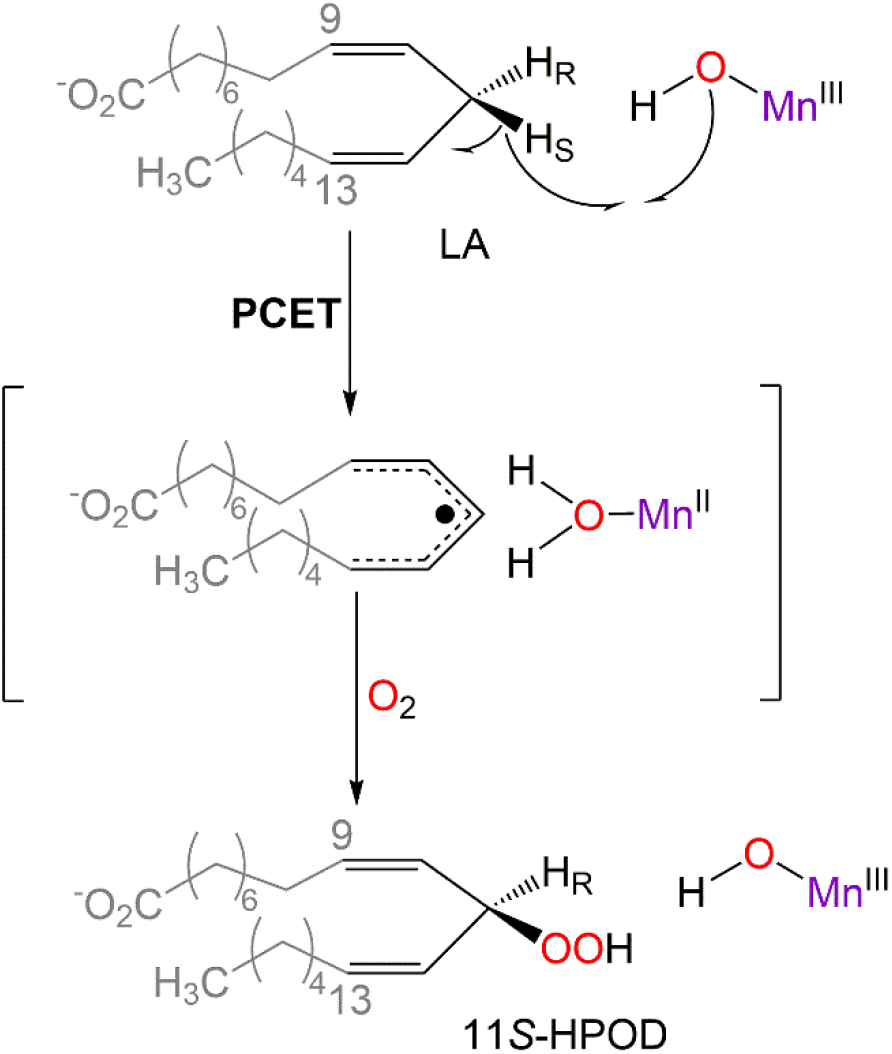
Mechanism of oxidation of linoleic acid, LA, by *Mo*LOX, involving a rate-limiting C-H cleavage catalytic step by proton-coupled electron transfer (PCET) involving a hydrogen tunneling mechanism.^*14*^ *Mo*LOX can also generate 9*S*/*R*- and 13*S*-hydroperoxides through β-fragmentation.^*13*^

As with most secreted enzymes, *Mo*LOX is a glycoprotein decorated with *N*-linked glycans on its surface. One of the inherent challenges in studying glycoproteins is the difficulty in characterization of the native protein structure because the branched and inherently flexible carbohydrates inhibit crystallization.^*15*^ To overcome this challenge, many studies resort to the modification or (partial or full) removal of the glycan(s) to enable crystal packing for high-resolution structural analysis. A crystal structure has been reported for *Mo*LOX with modified glycans and in the absence of a substrate at 2.04 Å resolution.^*16*^ This previous study used a *Mo*LOX sample treated with endonuclease H (EndoH), an endoglycosidase which removes mannose-rich oligosaccharides linked to asparagines on the surface of proteins (**Figure 1A**).^*16*^ However, for many *N*-linked glyco-enzymes, removal of the carbohydrates is linked to altered catalytic proficiency.^*17-20*^ As there is of yet no structure of the fully glycosylated form of *Mo*LOX, the impact of the removal of the *N*-linked glycans on the ES complex structure and on its enzymatic function is not well known. Therefore, to facilitate structure-guided inhibitor design for *Mo*LOX, it is imperative to resolve the structural details of the ES complex for the native glyco-enzyme.

**Figure 1.**
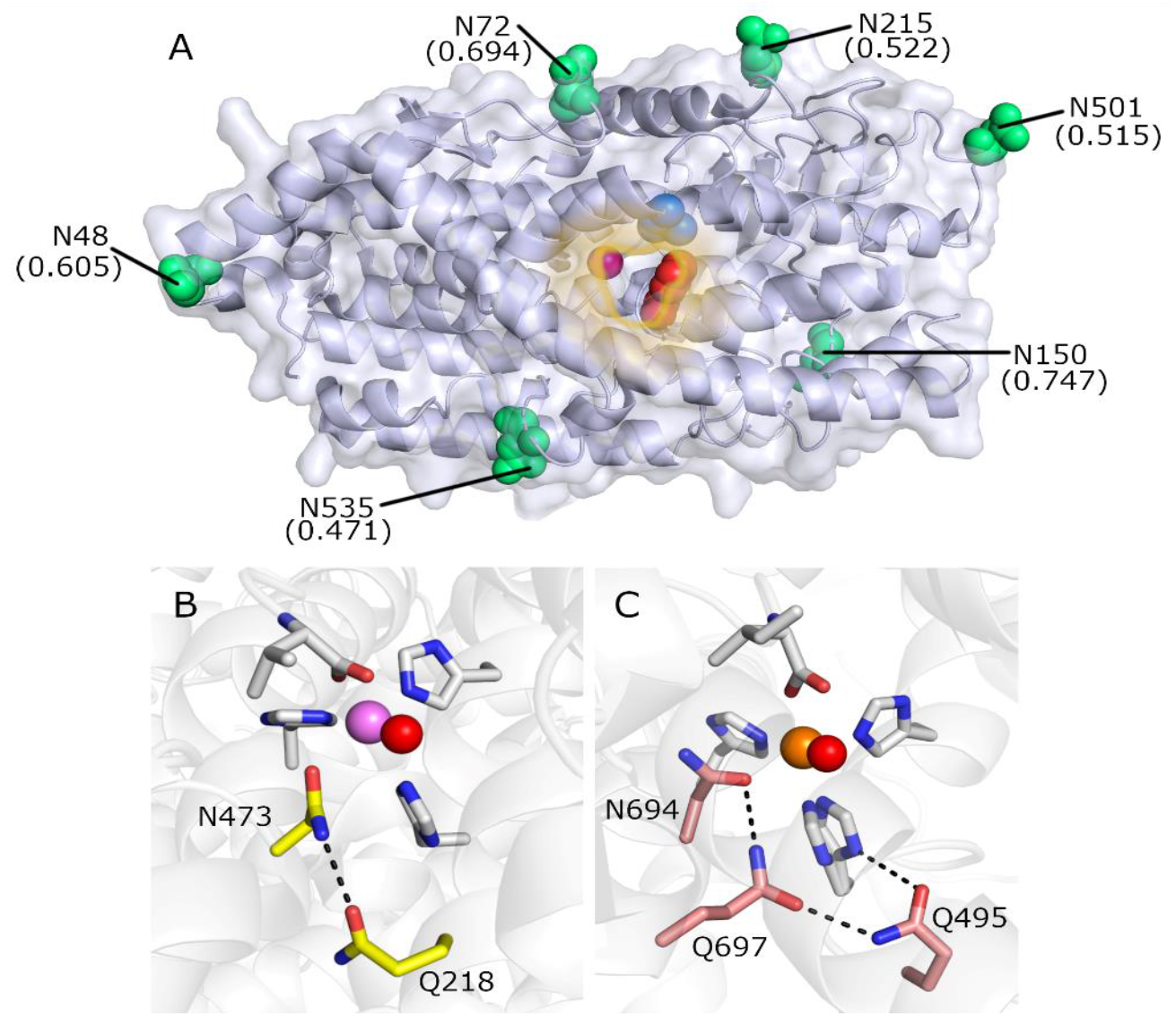
Structure^*16*^ of *Mo*LOX (A) with the predicted site Asn residues for *N*-linked glycosylation depicted in green spheres (N18 and N28 are missing from the X-ray structure). The parentheses represent the predicted potential for *N*-linked glycosylation. Also shown in spheres are the catalytic manganese center (purple) and the structurally conserved active site residues, L331 (red) and F526 (blue), that mediate substrate positioning for efficient hydrogen tunneling. (B) and (C) represent the first- and second-shell ligands to the mononuclear metal center in *Mo*LOX and soybean lipoxygenase (SLO), a model iron LOX. The second-shell ligands are labeled, and color coded for reference. Note that the *Mo*LOX numbering here is based on the processed enzyme that lacks the N-terminal signaling peptide sequence.

We previously utilized electron nuclear double resonance (ENDOR) spectroscopy to capture high-resolution, 3-D structural information about the elusive ES complex of the model plant lipoxygenase from soybean (SLO) with its substrate linoleic acid, LA.^*21, 22*^ Although ENDOR is most commonly used to characterize the coordination sphere of biological metal ions by interrogating ligands coordinated to the paramagnetic center,^*23*^ it can also provide high-precision structural information for atoms in the active site that are in close proximity to the metal ion but not covalently linked to it,^*24, 25*^ as is the case in the ES complex of SLO-LA.^*15*^ In the previous SLO study, to achieve the proper electronic properties for ENDOR analysis of the lipoxygenase the nonheme mononuclear iron was substituted by the non-native, high-spin (*S* = 5/2) Mn^2+^ ion as a spin-probe (Mn-SLO). ^13^C/^1^H ENDOR spectroscopy then revealed the position and orientation of substrate LA relative to the enzymatic metal center, and further gave information about the distribution in positioning.^*21, 22*^ ENDOR analysis and Molecular Dynamics (MD) simulation combined not only provide the important acceptor (Mn-Oxygen) to donor (LA-C11) distance (DAD) in the ground state, but also corroborate the previous theoretical understanding that, enzymes achieve reactive hydrogen tunneling geometries at the active site through protein thermal motions, starting from their ground state geometries.

Unlike the canonical Fe-LOXs found in prokaryotes, plants and animals, LOXs found in pathogenic fungi, such as *Mo*LOX, are naturally equipped with a catalytically active Mn^2+^ cofactor (Mn-LOX).^*13, 26, 27*^ In the current study, we use EPR and ENDOR spectroscopies to obtain the first ground state active-site ES structure for the native, *N*-linked glycosylated form of the fungal Mn-LOX, *Mo*LOX, and compare that to the ES structure of a de-glycosylated form of *Mo*LOX to test the effects of the surface glycans on the active-site structure. Simulation of the Mn^2+^ EPR spectrum of *Mo*LOX is consistent with the subtle, yet catalytically important, differences of the first- and second-shell ligands surrounding the mononuclear metallocentre of Fe-SLO and *Mo*LOX, seen in their respective X-ray crystal structures (**Figure 1B, C**). Most importantly, the high-spin Mn^2+^ ion of *Mo*LOX acts as a spin probe, enabling the sensitive use of ^13^C ENDOR to determine the positioning of substrate LA relative to the *catalytically active* metal ion in the ES complex of *Mo*LOX. The ENDOR results for the native *Mo*LOX-LA complex are further compared to those of a de-glycosylated form of *Mo*LOX prepared by treatment with EndoH, and this comparison is complemented with comparison of kinetic isotope effects that report on the effects of glycan processing on enzyme activity. Finally, the ENDOR-determined Mn^2+^-LA distances are used as restraints for MD simulations of the ES complex that yield the first high-resolution image of the complete active site of the native *Mo*LOX-LA co-structure, in particular giving the distance from the oxygen-atom bound to Mn that accepts an H-atom from the reactive carbon donor of substrate (**Scheme 1**; C11↔O). The results provide unique insights into structural and functional differences across the evolutionarily divergent family of Fe- and Mn-LOXs.

## Results

*Mn*^*2+*^ *EPR*: We start by describing the Mn^2+^-EPR spectrum of native *Mo*LOX in the light of protocols established earlier for the manganese-substituted soybean lipoxygenase (Mn-SLO).^*21*^ The EPR spectrum of the *S* = 5/2 Mn^2+^ ion of *Mo*LOX (**Figure 2a**), like that of Mn-SLO (**Figure S1**), is the sum of the ‘envelopes’ of the five orientation-dependent Δ*m*_*s*_ = ± 1 transitions between adjacent pairs of the six electron-spin m_s_ substates, -5/2 ≤ *m*_*s*_ ≤ +5/2. Each envelope has a well-defined shape that is spread over a range of fields defined by its *m*_*s*_ values and the magnitudes of the zero-field splitting (ZFS) parameter, *D*, and a rhombicity parameter, λ, with each envelope weighted by the thermal population of the contributing m_*s*_ levels. The overall breadth of the pattern is proportional to the magnitude of *D*, which in turn increases with deviations of the coordination sphere of the Mn^2+^ ion from spherical symmetry; details of the shape of the pattern are determined by the rhombicity parameter, 0 ≤ λ ≤ 1/3, whose value increases with deviations from axial symmetry around the principal ZFS axis (λ = 0).

**Figure 2:**
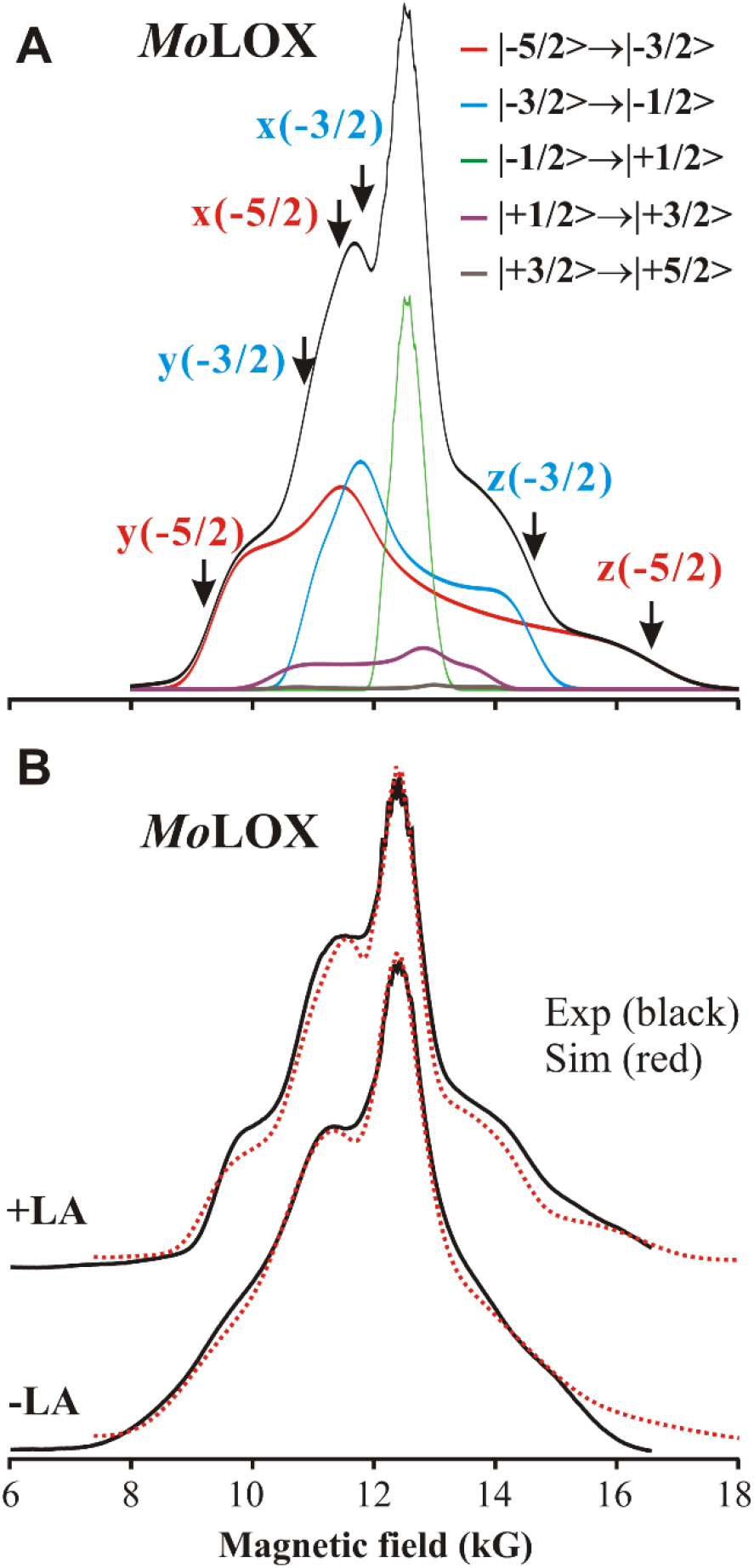
(upper) *Mo*LOX EPR spectra: absorprion-display EPR envelopes of individual contributions from five electronic transitions as differentiated by colored lines. (lower) 35 GHz ESE-EPR spectra of *Mo*LOX with and without LA. Simulation in red. Exp conditions: MW freq 34.9 GHz, MW pulse length (π/2) = 40 ns, τ = 500 ns, repetition rate =100 Hz, Temperature 2 K, Simulation parameters see Table1.

The value of *D* of *Mo*LOX is three-fold larger than that of Mn-SLO (**Figure 2b, Figure S1**, see **Table 1** for *Mo*LOX *D* = 3500 MHz,) which is manifested as three-fold larger magnetic field span of the EPR spectrum of *Mo*LOX than that of Mn-SLO. The Mn-coordination sphere of *Mo*LOX is a distorted octahedron that doesn’t overlap exactly with the Fe-SLO structure (**Figure 1b**,**c**).^*16*^ This subtle difference in the position and orientation of Mn-coordinating ligands in the two Mn enzymes causes the appreciable differences in ZFS as evidenced in the EPR spectra. Another native, functional Mn-LOX from the fungal family *Gaeumannomyces graminis, Gg*LOX also shows a broad EPR spectrum like that of *Mo*LOX and has similar ZFS parameter (**Fig. S1, Table 1**).^*27*^

**Table 1.** ZFS Parameters of Mn^2+^ in Mn-LOX

| Mn-LOX | substrate | $D$ (MHz)<sup>a</sup> | $\Lambda$ | $f$ |
| --- | --- | --- | --- | --- |
| *Mo*LOX | -LA | +3500 | 0.14 | 0.6 |
|  | +LA | +3000 | 0.17 | 0.3 |
| *Gg*LOX<sup>b</sup> | -LA<sup>(i)</sup> | +2100 to +3000 | 0.13-0.23 | N.R. |
| Mn-SLO | -LA/+LA | +1200 | 0.17 | 0.3 |

a) $D$ is the axial ZFS parameter, $\Lambda = E/D$ is the rhombicity. In the simulations, $D$ and $E$ each has a width that is the fraction, $f$ , times the parameter (See SI).
b) Native manganese lipoxygenase from fungus, *G. graminis*; values reported from ref <sup>27</sup> N.R., not reported.

Unlike Mn-SLO, the EPR spectrum of *Mo*LOX is notably changed by the addition of the substrate, LA, (**Figure 2b** and **S1**) with well-defined sharpening of the contribution from the different electron spin manifolds, corresponding to a two-fold decrease in the distribution in the ZFS parameters, along with a small reduction in the ZFS splitting, *D* (**Table 1**). The distribution in ZFS parameters reflects the degree of flexibility to the Mn coordination sphere. Although LA is not directly ligated to Mn^2+^and rather is in the first coordination sphere as we will learn from ENDOR analysis (see below), the sharpened Mn^2+^ EPR spectrum upon substrate binding indicates improved ordering of the Mn coordination geometry, while the small reduction of ZFS splitting suggests a subtle change in the ligand arrangement.

*ENDOR of Mn*^*2+*^ *(S = 5/2) in MoLOX*. For a single molecular orientation of a paramagnetic center of spin ***S=5/2***, the first-order ENDOR spectrum for an ***I*** = 1/2 nucleus (^13^C, ^1^H) obtained by monitoring an EPR transition between adjacent substates, *m*_*s*_ ⇔*m*_*s*_ +1 comprises a signal from each of the substates. For this study, the two frequencies are conveniently written as offsets (δν_*ms*_) from the nuclear Larmor frequency, ν_N_,

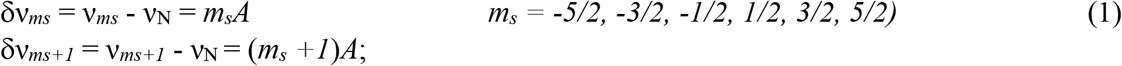

where *A* is the orientation-dependent electron-nuclear hyperfine coupling interaction.

A ^13^C of the non-coordinated, labelled LA substrate experiences a through-space dipolar interaction between the electron spin of the Mn^2+^ center and the ^13^C nuclear spin. Although the Mn^2+^ spin is partially delocalized over its coordinating ligands, for the interaction with a relatively remote ^13^C of bound LA this hyperfine interaction can be taken as a point dipole coupling with the Mn^2+^ ion, **T**(*r*),

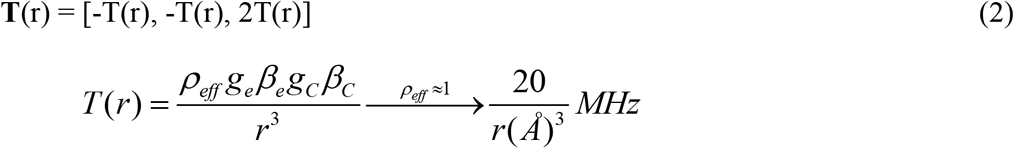

whose magnitude is determined by the length of the Mn^2+^ → ^13^C vector (*r*), and whose unique axis points along this vector; it incorporates an effective Mn spin density that can be taken as, ρ_eff_ ≈ 1.^*21*^ As described in the Methods section, simulation of the full 2D field-frequency pattern of multiple ENDOR spectra collected across the entire EPR envelope of *Mo*LOX-LA, as exemplified by the experimental 2D patterns for ^13^C10 and ^13^C11 of LA bound to *Mo*LOX **Fig 3**, enables determination of the dipolar interaction tensor, ***T***, for each of the two nuclei ^13^C (C10, C11), thereby yielding the Mn-C distances *r* **(eq 2**) and the orientation of the two Mn-C vectors relative to the ZFS coordinate frame for the Mn^2+^ ion.

**Figure 3:**
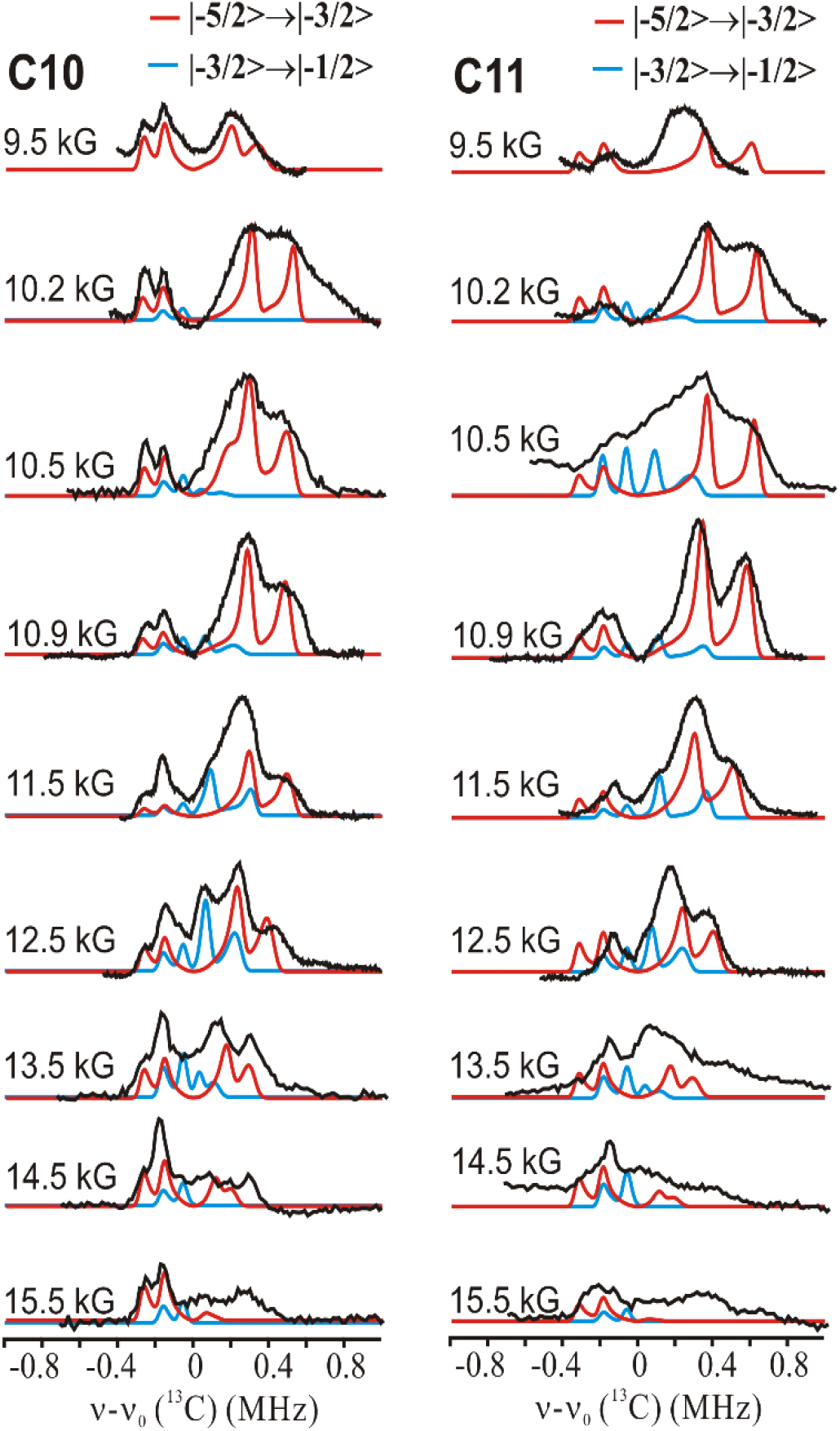
35 GHz 2D field-frequency pattern ^13^C Mims ENDOR for *Mo*LOX WT with ^13^C10-LA and ^13^C11-LA and simulations that highlight peak positions, not shapes (in red and blue ENDOR contributions from two electron-spin transitions as mentioned; simulation parameters, **Table 2**). Conditions: microwave frequency ∼ 34.8 GHz, MW pulse length (π/2) = 50 ns, τ = 1500 ns, repetition rate =100 Hz, temperature 2 K.

The analysis of the 2D patterns of ^13^C10 and ^13^C11 of LA bound to *Mo*LOX, **Figure 3**, began by noting the overall similarities of these patterns to the corresponding patterns previously seen for LA bound to Mn-SLO.^*21*^ In SLO, C10 and C11 of LA lie in the x-z ZFS plane (Euler angle φ = 0), and so at the low-field edge of the EPR spectrum, where the magnetic field pointed along y, the magnetic field is perpendicular to a Mn-C vector, and the ENDOR spectrum shows a doublet whose splitting is precisely T(r), whose value yields the Mn-C distances directly (**eq 2**); the full analysis of the 2D pattern then determined the remaining Euler angles, θ.

**Table 2.** ENDOR hyperfine parameters for ^13^C-LA.

| | T(MHz) | r(Å) | ($\theta$, $\varphi$) |
| --- | --- | --- | --- |
| *MoLOX* |  |  |  |
| C10 | 0.11 | 5.7 | 90, 15 |
| C11 | 0.13 | 5.3 | 90, 30 |
| H<sub>2</sub>O | 4.2 | 2.7 | 90, 0 |
| *Mn-SLO* |  |  |  |
| C10 | 0.17 | 4.8 | 50, 0 |
| C11 | 0.17 | 4.8 | 40, 0 |
| H<sub>2</sub>O | 4.0 | 2.7 | 70, 0 |

Comparison of the 2D ^13^C ENDOR patterns of **Fig 3** with 2D patterns calculated for trial Mn-C orientations chosen from a grid of (θ, ϑ) values showed that the Mn-C vectors of C10 and C11 of *Mo*LOX both lie in the ZFS (x, y) plane (θ = 90°). However, φ ≠0, in which case the splittings in the low-field ENDOR spectra do not simply yield T(r). To determine T(r) we used the fact that as the field is varied across the EPR envelope, the maximum splitting of the ^13^C ENDOR doublet observed at frequencies greater than the ^13^C Larmor frequency (**Fig 3**) is associated with the subset of orientations with the external field parallel to the Mn-C vector, and that this maximum splitting equals 2T(r). This approach yields T(r) = 0.11 MHz for ^13^C10 and 0.13 MHz for ^13^C11 (**Figure 4**), corresponding to Mn-C distances of *r* = 5.7 Å for C10 and 5.3 Å for C11. Although, the reactive C11 carbon of substrate LA in *Mo*LOX is significantly farther from the metal center than in Mn-SLO (Mn-C11 = 4.8 Å), as we show below, when using these distances as constraints on MD calculations, the distances between C11 and the reactive O bound to Mn are found to be nearly the same in the two enzymes.

**Figure 4:**
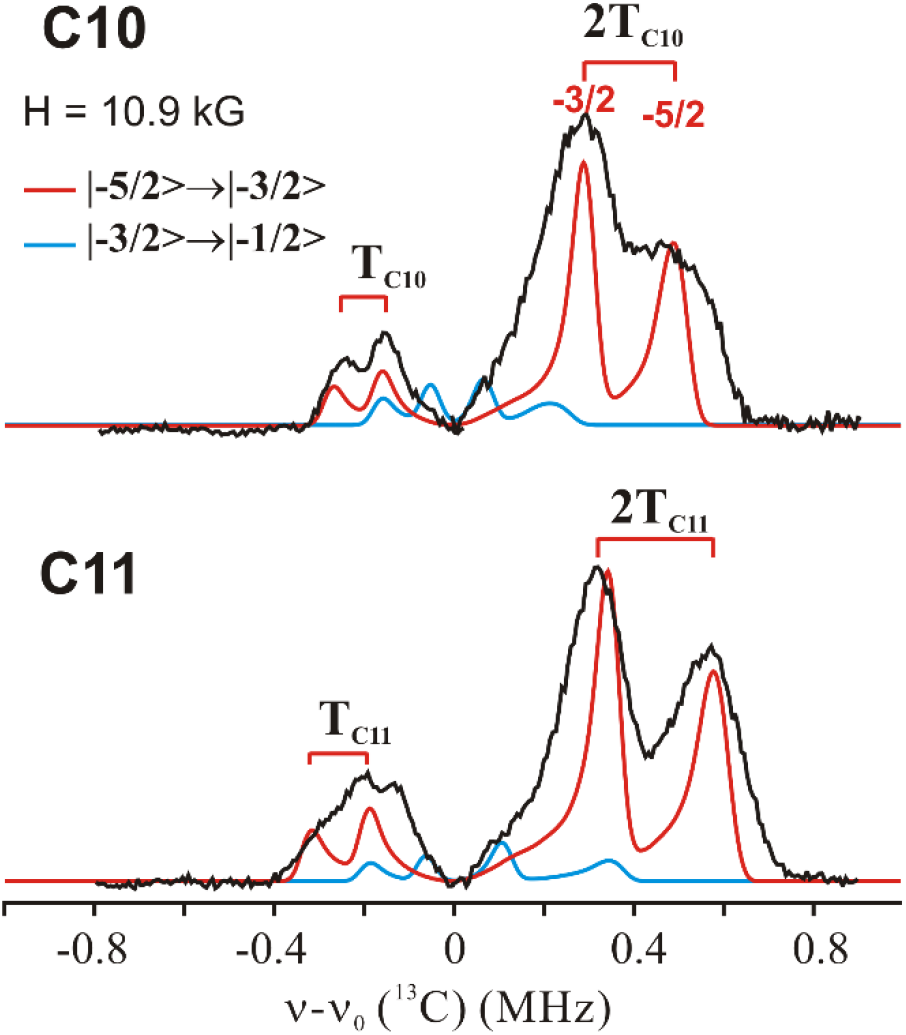
35 GHz ^13^C Mims ENDOR of WT *Mo*LOX with ^13^C10-LA, ^13^C11-LA. Simulation (in red and blue) of the ENDOR contributions from two electron-spin transitions as mentioned. Conditions are same as in **Figure 3**.

The values for T(r) and θ = 90° for C10 and C11 are well determined, as is φ ≈ 15° for C10. However, uncertainties in the optimum values of φ for C11 led us to carry out an additional joint optimization, as follows. Taking the Mn-C distances given above for ^13^C10 and ^13^C11, along with a C10-C11 bond distance of 1.55 Å, the law of cosines gives the angle between the Mn-C vectors, δφ ≈ 15-16°. Then, as both vectors were found to lie in the ZFS xy plane (θ = 90°), the well determined value φ ≈ 15° for C10, then leads to φ ≈ 30° for C11. The resulting simulations of the 2D ENDOR patterns for the two carbons are shown in **Figure 3**. As the simulations show, the ^13^C ENDOR spectra of C10, C11 (**Figures 3** and **4**) are dominated by responses from -5/2 to -3/2 electronic transition (in red). However, the ENDOR contribution from -3/2 to -1/2 transition, though weak, is needed to match additional peaks in the ENDOR pattern.

The ^13^C10 and ^13^C11 ENDOR spectra of *Mo*LOX-LA both display broader linewidths with less resolution than those from Mn-SLO (**Figure S2**), which can be understood considering the following two points. (1) Although *Mo*LOX has a larger ZFS (**Table 1**) which improves orientation selection, it also has a bigger dispersion/distribution in *D*, which more strongly acts to diminish orientation selection. (2) The different orientations of the Mn-C vector relative to the ZFS coordinate frame in *Mo*LOX means that in the low-field spectra the field does not lie normal to the Mn-C bond and sample a distribution in hyperfine couplings.

Interestingly, the ^13^C ENDOR of Mn-SLO revealed a longer Mn-C11 distance, 5.7 Å, assigned to an inactive ‘*b*’ state. If such a second conformer with a longer Mn-C11 distance does exist in *Mo*LOX, then the resulting, even smaller dipolar interaction would give ENDOR features buried around the ^13^C Larmor frequency and thus could not be detected.

### ^*1*^H_2_O ENDOR of MoLOX-LA

To orient the C10-C11 fragment of LA relative to the Mn-OH_2_ linkage, and thereby generate the information necessary to estimate the metal-bound oxygen-to-carbon distance central to catalytic H atom abstraction, we carried out ^1^H Davies ENDOR measurements on the exchangeable protons of the metal-bound H_2_O (**Figure S3**), isolating the ^1^H signals from bound H_2_O signals from those of the constitutive protons of coordinated histidine by subtracting spectra collected with enzyme in D_2_O buffer from those of enzyme in H_2_O buffer. Simulation of the resulting ^1^H 2D ENDOR pattern (**Figure 5**) shows that the Mn-^1^H vector lies along the ZFS x-axis (θ, ϑ) = (90, 0). As a result, at the low (H = 9.5 kG) and high (H = 16.5 kG) field edges of the EPR spectrum, ^1^H ENDOR yields single crystal-like spectra; the low-field ^1^H doublet splitting is T = 4.2 MHz, which corresponds to a Mn-H distance of 2.7 Å for a through-space Mn-H dipolar interaction. There is no detectable response from a second water proton. Taking the Mn-O distance of 2.23 Å and an Mn-O-H angle of ∼ 109^°^, simple geometric considerations allow a range of possible dihedral angles between the Mn-O-H and Mn-O-C11 planes, in turn allowing an O↔C11 distance that can be as short as ∼ 3.1 Å. Nonetheless, the imposition of the constraints associated with the Mn-O^1^H measurement is of importance in the MD simulations described below.

**Figure 5:**
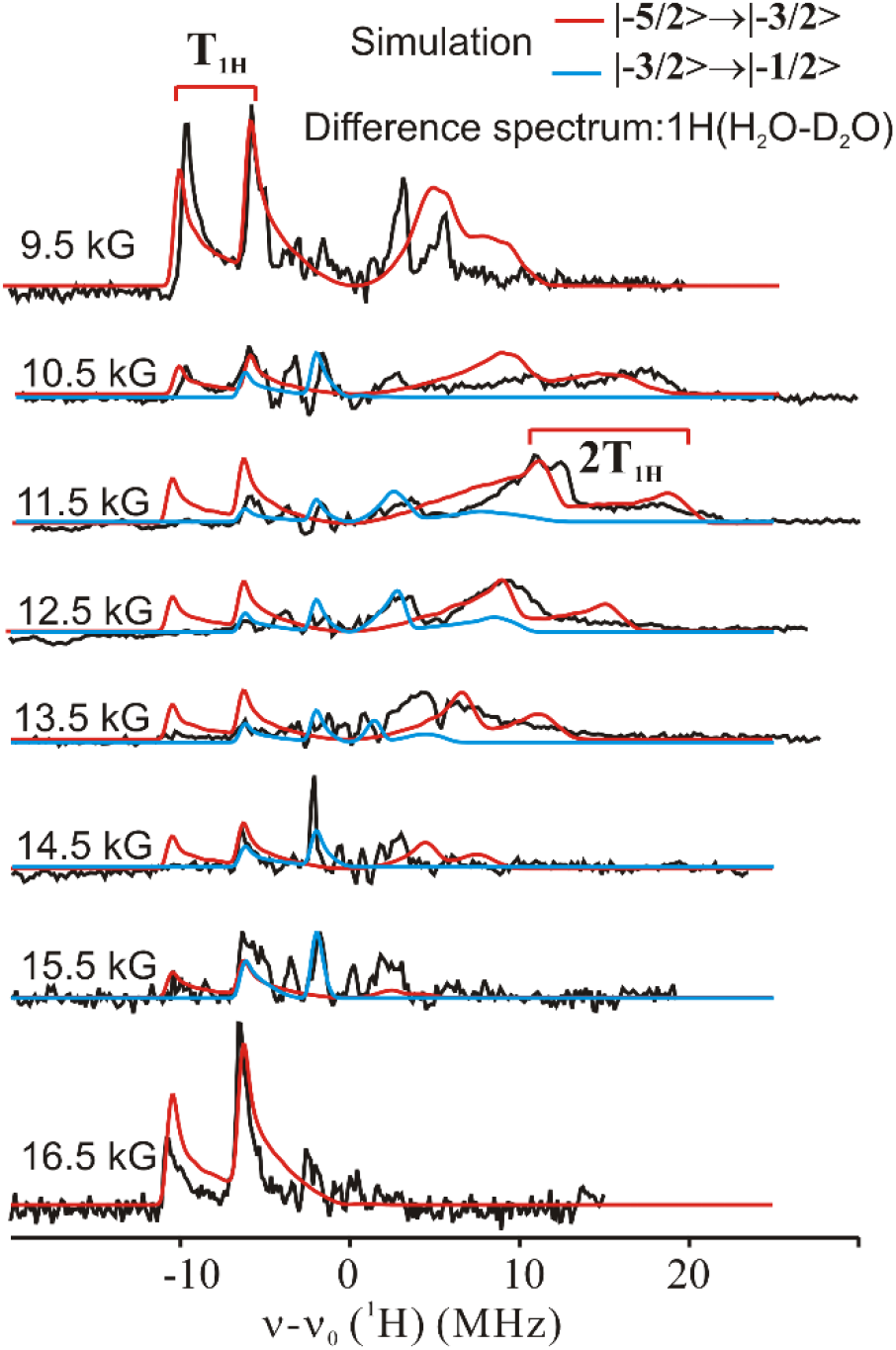
35 GHz 2D field-frequency pattern ^1^H Davies ENDOR for WT *Mo*LOX with LA and simulation (in red and blue) ENDOR contributions from two electron-spin transitions as mentioned. Conditions: microwave frequency ∼ 34.8 GHz, MW pulse length (π/2) = 50 ns, τ = 500 ns, repetition rate =100 Hz, temperature 2 K. Simulation parameters are listed in Table 2.

### Studies of EndoH Modified MoLOX-LA

The published X-ray structure of *Mo*LOX (**Figure 1A**)^*16*^ was collected from a modified form of the protein in which the enzyme was pre-treated with endonuclease H (EndoH), an endoglycosidase which removes mannose-rich oligosaccharides linked to asparagines on the surface of proteins, leaving an *N*-GlcNAc attached. Wild-type (WT) *Mo*LOX has seven predicted sites for *N*-linked glycans.^*14*^ Removal of the *N*-glycans by EndoH reduces the molecular mass of the protein from 110-130 kDa in the native state to ∼70kDa, as shown in our SDS-PAGE analysis (**Figure S4**) and in agreement with the literature.^*13*^ Such a change in molecular mass raises the question as to what degree could removing the *N*-glycans from the *Mo*LOX surface influence the active site structure and enzyme function. To validate the use of the EndoH-modified X-ray structure of *Mo*LOX for MD simulations, we also examined the ^13^C ENDOR responses for this de-glycosylated sample.

### ^*13*^C ENDOR of EndoH-Treated MoLOX

The EPR spectrum of EndoH-treated *Mo*LOX is identical to WT both in the presence and absence of substrate LA, which suggests that the local Mn^2+^ coordination environment is unaffected by the surface modifications (**Figure S5**). The field dependence of the ^13^C Mims ENDOR spectra of ^13^C10, ^13^C11 labeled LA of EndoH-treated *Mo*LOX-LA matches nicely with that of WT (**Fig S5**). The ENDOR peak positions and linewidth are essentially unchanged, with the spectra completely overlaying those of WT (**Figure 6** and **S6**). The unchanging peak frequencies show that the orientation and positioning of the substrate doesn’t change upon clipping of glycans, while the unaffected linewidths indicate that the precision with which the active site positions the substrate also does not change. In short, the ENDOR measurements indicate the ground state active site structure of the ES complex is unchanged by the clipping of glycans by EndoH.

**Figure 6:**
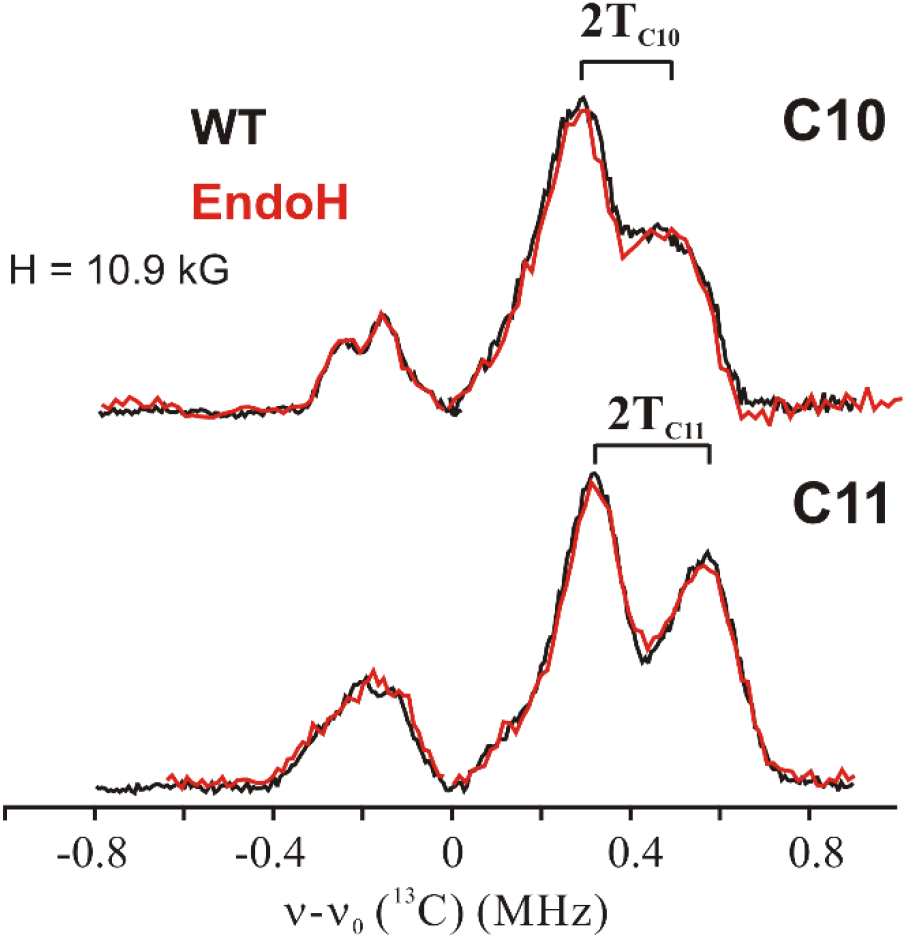
35 GHz ^13^C Mims ENDOR of native (***black***) and EndoH (***red***) *Mo*LOX with ^13^C10-LA, ^13^C11-LA. ENDOR contributions from two electron-spin transitions as mentioned. Conditions same as in **Figure 3**. H = 10.9 kG. The complete 2D ENDOR spectra are in **Figure S6**.

### Kinetic Studies of WT and EndoH MoLOX

The ENDOR-derived invariant positioning of LA in WT and EndoH-treated *Mo*LOX is consistent with the analysis of its steady-state kinetic parameters (**Table 3**). Primary deuterium isotope effects (^D^*k*_cat_) and their temperature dependence, Δ*E*_a_, (**Table 3**) report on the proficiency of substrate positioning and dynamics at the active site for enzymatic C-H reactions.^*28*^ The Δ*E*_a_ of the EndoH *Mo*LOX variant (−0.9 ± 1.5 kcal/mol) was, within uncertainty, the same as that of the native, glycosylated enzyme, with both being close to ∼0, supporting a hydrogen tunneling mechanism. The unchanged Δ*E*_a_ following EndoH processing demonstrates that the loss of surface glycans does not alter the properties of tunneling associated with catalysis. Taken together, the kinetic and ENDOR observations indicate the substrate positioning in the active site is not significantly perturbed when the *N*-glycans are removed from the surface of *Mo*LOX by EndoH.

**Table 3.** Kinetics of native and EndoH-treated *Mo*LOX

|  | WT *MoLOX* | EndoH *MoLOX* |
| --- | --- | --- |
| $k_{\text{cat}}$ ($\text{s}^{-1}$)<sup>a</sup> | $2.41 \pm 0.17$ | $2.38 \pm 0.15$ |
| $k_{\text{cat}}/K_{\text{M}}$ ($\mu\text{M}^{-1} \text{s}^{-1}$)<sup>a</sup> | $0.17 \pm 0.03$ | $0.12 \pm 0.02$ |
| $E_{\text{a}}(\text{H})$ (kcal/mol) | $9.5 \pm 0.5$ | $8.5 \pm 1$ |
| $^{\text{D}}k_{\text{cat}}$ <sup>a,b</sup> | $70 \pm 2$ | $70 \pm 6$ |
| $\Delta E_{\text{a}}$ (kcal/mol)<sup>c</sup> | $-1 \pm 0.6$ | $-0.9 \pm 1.5$ |

\*In borate buffer, pH 9.0 @ 20°C. <sup>bD</sup> $k_{\text{cat}} = k_{\text{cat}}(\text{H})/k_{\text{cat}}(\text{D})$ . <sup>c</sup> $\Delta E_{\text{a}} = E_{\text{a}}(\text{D}) - E_{\text{a}}(\text{H})$ .

### MD Simulated Model of MoLOX-LA Complex

The final model of the LA structure in *Mo*LOX was generated by MD simulations that used the X-ray crystal structure of EndoH-*Mo*LOX to obtain the starting coordinates and by then imposing the C11−Mn and C10−Mn ENDOR-derived distances as constraints. Forty ns of MD trajectories, treated as two independent 20 ns trajectories, were carried out for the LA-EndoH-*Mo*LOX complex using harmonic restraints from the ENDOR-derived Mn-C11 and Mn-C10 distances. The final snapshot of the second 20 ns trajectory is presented in **Fig 7A**. In this model, LA is positioned within the U-shaped substrate channel in the *Mo*LOX active site proximal to the catalytic Mn center (shown in yellow and purple sticks). The carboxylate moiety of LA is located near the surface and is predicted to hydrogen bond with Arg525 and Arg528. This “carboxylate out” binding configuration is consistent with that predicted from a published docked model (**Fig. S8**),^*16*^ and also is consistent with the co-structure of the coral 8*R*-LOX with arachidonic acid.^*8*^ The ensemble average distance from the C11 hydrogen donor to the Mn-bound oxygen acceptor (i.e., DAD; **Fig 7B**) obtained from these MD simulations is 3.4 ± 0.3 Å (**Fig 7C**). This DAD is approximately the sum of the van der Waals radii for carbon and oxygen, *ca*. 3.2 Å.

**Figure 7.**
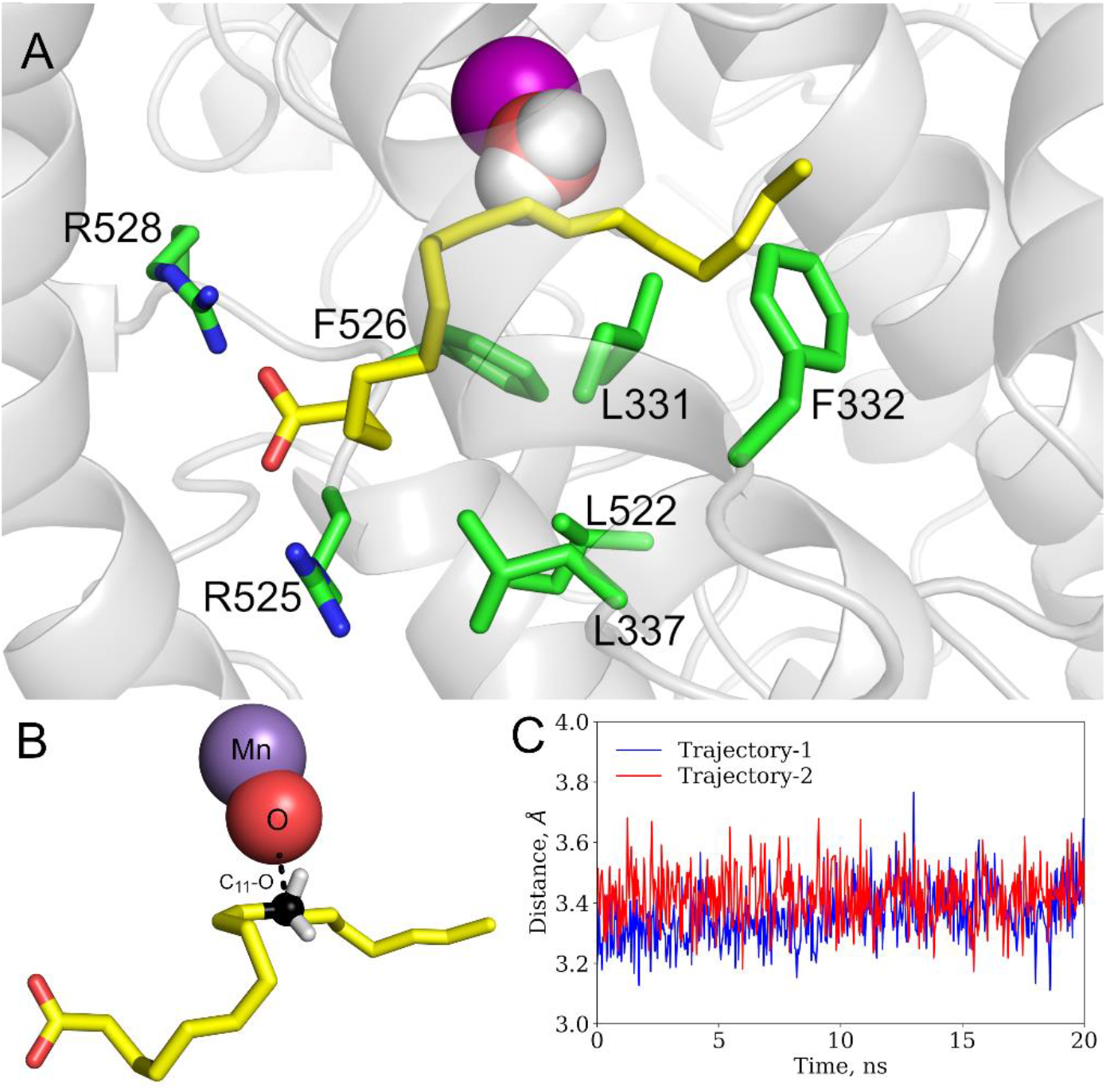
MD model of LA bound to active site of *Mo*LOX. (A) MD snapshot of the *Mo*LOX complex with LA (final snapshot). For comparison, overlays of the *in silico* docked model and snapshots of the two MD trajectories are shown in **Figure S8**. The substrate is represented as yellow sticks. Potentially important sidechains involved in substrate positioning are labeled and represented as green sticks. (B) Representation of the DA distance between carbon-11 (black) of substrate and Mn-bound oxygen (from water). (C) C11↔O distance along the two 20 ns MD trajectories for LA in native *Mo*LOX (*cf*. **Fig S9**).

### Kinetic Studies of *Mo*LOX Active Site Mutants

To validate the MD model in **Figure 7A**, we prepared a series of enzyme variants in which five aliphatic residues predicted to line the substrate-binding site (i.e., within 4 Å of LA): L331, F332, L337, L522, and F526, were individually mutated to alanine and tested the kinetic impact (**Table 4**). Similar strategies have been employed to map the important residues involved in substrate binding and positioning in other lipoxygenases.^*30-32*^ For example, alanine substitution of conserved Leu residues in SLO and human 12-LOX have been reported to decrease the rate constant *k*_cat_ by 100- to 1000-fold.^*32, 33*^. The alanine substitutions at the homologous residues in *Mo*LOX, L331 and F526, which sit within 4 Å of C11 of LA, cause an ∼20-fold decrease in *k*_cat_, which confirms the expectation that these residues aid in positioning C11 of LA close to the metal-bound oxygen in *Mo*LOX.

**Table 4.**
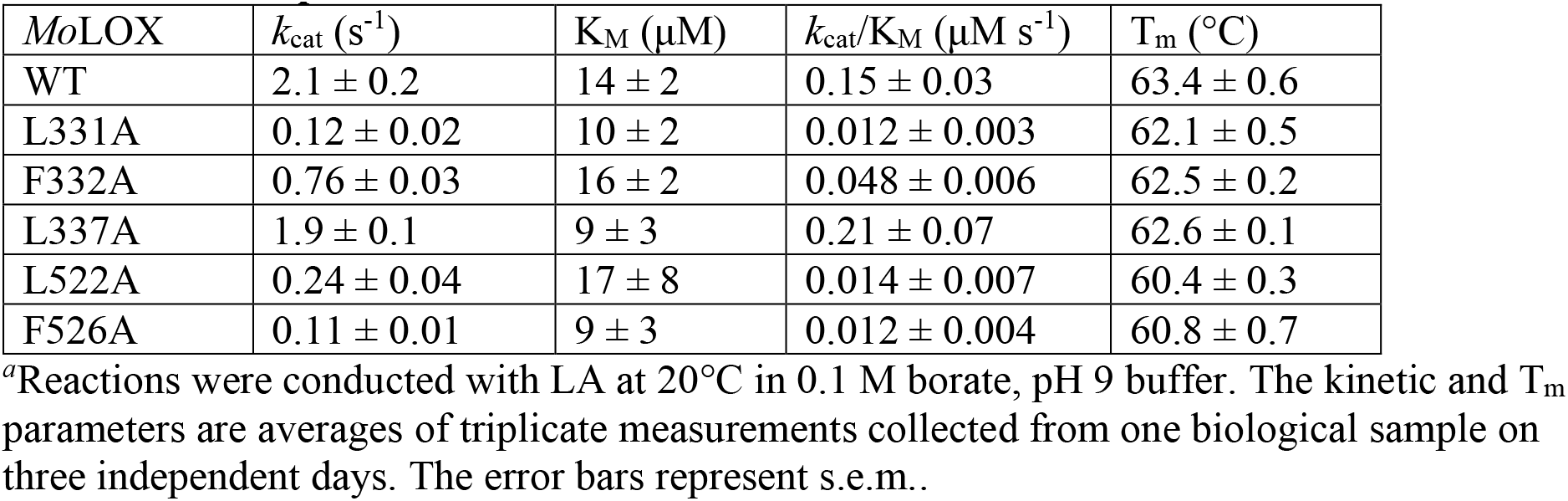
Kinetic parameters of *Mo*LOX active site mutants^*a*^.

In the *Mo*LOX-LA model, F332 is within 4 Å of C15 and C16 of LA and adjacent to L331 (see above). Substitution of Phe-for-Ala results in a modest 3-fold decrease in *k*_cat_ (**Table 4**). It is important to note that F332 is located in a loop that undergoes significant movement during the 40 ns MD trajectory. The Cα of F332 in the final snapshot of the MD simulation (**Figure 7**) is displaced by 3 Å relative to the Cα position in the X-ray structure (**Figure S8**). To probe if these dynamics have any relevance to catalysis, we also measured the activation energy (*E*_a_) for F332A. In the context of hydrogen tunneling, changes in the *E*_a_ value are consistent with altered *isotope-independent* conformational (thermal) motions.^*11, 29*^ The *E*_a_ of F332A is reduced (7.2 ± 0.7 kcal/mol) relative to WT *Mo*LOX, *E*_a_ = 9.5 kcal/mol (**Table S1** and **Figure S7**), supporting a functional role for the dynamic nature of this loop. The kinetic behavior of F332A *Mo*LOX is reminiscent of the SLO I553X mutants (X = L or G) that showed little-to-modest changes in *k*_cat_, but was accompanied by altered *E*_a_ values that are associated with disruption of active site sidechain packing and a network of protein motions.^*34-36*^

Finally, we explored alanine mutations at L337 and L522 in *Mo*LOX (**Table 4**). Kinetic analysis of L337A and L552A show distinct catalytic impacts in *Mo*LOX compared to their homologous mutants, I553A and V750A,^*36*^ in SLO (**Table S1**). Mutation to alanine of L522, which is positioned behind and contacts both L331 and F526, produces a 9-fold decrease in *k*_cat_. This demonstrates the importance of proper hydrophobic packing in the *Mo*LOX active site. In contrast, the L337A *Mo*LOX variant has a nearly identical *k*_cat_ to that of WT. Further, its *E*_a_ of 9.9 ± 1 kcal/mol is unchanged from that of WT (9.5 kcal/mol; **Figure S7**). Conversely, the homologous mutation in SLO, V750A, exhibits a decreased *E*_a_ with a slightly reduced *k*_cat_ value relative to WT. Note that the second-order rate constants, *k*_cat_/K_m_, for this suite of *Mo*LOX mutants follow similar trends to *k*_cat_ (**Table 4**). The cumulative kinetic properties validate the LA-EndoH-*Mo*LOX binding model in **Figure 7A** while the differences from the kinetic properties of SLO illustrate the influence of subtle differences in substrate positioning by aliphatic, active-site residues.

## Discussion

### ENDOR discussion: EPR/ENDOR analysis of Mn-SLO vs MoLOX

We previously used EPR/ENDOR spectroscopy to identify the enzyme-substrate (ES) ground state structure in metal substituted (Mn^2+^-for-Fe^2+^) soybean lipoxygenase, SLO.^*21, 22*^ Here we report the first EPR/ENDOR study of the ES complex of a fungal Mn-lipoxygenase, *Mo*LOX, which did not require metal substitution. Measurement of the ^13^C hyperfine coupling of C10 and C11 nuclei of LA, yielded the position and orientation of LA at the active site. Incorporation of this information as constraints on MD simulations that used the EndoH-*Mo*LOX crystal structure as starting coordinates has yielded the first high-resolution image of the active site of the *Mo*LOX-LA co-structure, the first for a fungal lipoxygenase in its native, fully *N*-glycosylated state, **Figure 7A**.

To begin we compare the structural information derived from EPR/ENDOR results for ES complex of *Mo*LOX-LA in the light of canonical Mn-SLO-LA. The EPR spectrum of Mn^2+^ at the enzymatic site provides information about the symmetry of the coordinating ligands. Mn(II)-*Mo*LOX shows a broader EPR spectrum than for Mn(II)-SLO as the result of a larger ZFS splitting, which suggests a greater distortion of the Mn-coordination sphere from octahedral symmetry for *Mo*LOX than Mn-SLO. This reflects the fact that the structures of the Mn-cofactor in the two enzymes as revealed by X-ray diffraction are not identical and do not overlap (**Fig 1b, 1c**), with small but distinct differences in orientation and position of metal ligands. These subtle structural differences in the metal geometry/symmetry likely account for the functional differences between the active Mn-containing *Mo*LOX and inactive Mn-substituted SLO.

The most obvious difference between the ^13^C ENDOR-derived LA positioning in Mn-SLO and *Mo*LOX is the distance between Mn and the C11 target, which is significantly greater in *Mo*LOX, 5.3 Å, than in Mn-SLO, 4.9 Å (**Figure 8**). Nonetheless, considerations of the ^1^H ENDOR of Mn-bound H_2_O show the distance between the aqua oxygen and target C11 can be as short as ∼ 3.1 Å for *Mo*LOX, the same as the estimated average ground-state distance of 3.1 Å in Mn-SLO. Note that ENDOR analysis of Mn-SLO led to the identification of two LA-conformers distinguished by Mn-LA distance (active conformer Mn-C11 4.9 Å, inactive conformer Mn-C11 5.7 Å), while there appears to be only single dominant LA conformer in the active site of *Mo*LOX.

**Figure 8:**
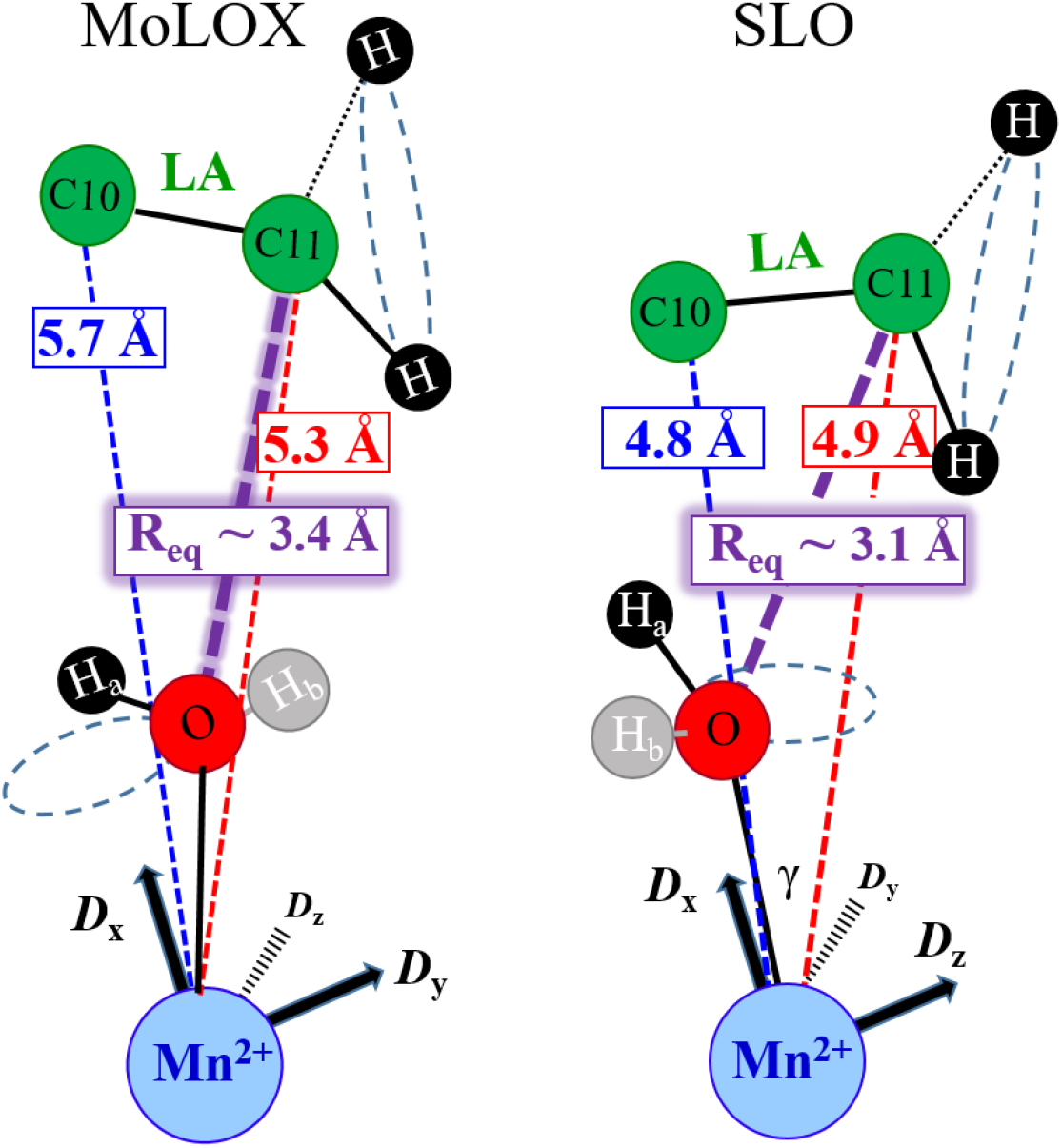
ENDOR-derived structure of active site of native *Mo*LOX-LA (left) and SLO-LA active conformer (right). SLO structure was adapted with permission from ref 21. Copyright 2017 American Chemical Society.

In short, although ENDOR measurements show the Mn-C11 distances differ in the two enzymes, differences in the organization of aliphatic sidechains along the substrate binding channel, nonetheless allow the *Mo*LOX ES complex to access comparable values for the catalytically important O↔C11 donor-acceptor distance (DAD). Of particular note, the ENDOR-derived distances are in excellent agreement with the MD-derived DAD of 3.4 (±0.3) Å.

The ENDOR-guided MD active-site structure of the ES complex described above was initiated using a docked model of LA into the available X-ray structure of EndoH-treated *Mo*LOX. Importantly, the ^13^C10, ^13^C11 dipolar interactions remain unchanged after clipping glycans (EndoH-*Mo*LOX), which indicates that the substrate LA positioning and orientation is not perturbed in the active site upon de-glycosylation. This observation, taken together with the kinetic impact of the volume-reducing alanine mutational screening of a series of aliphatic residues that line the putative substrate channel in *Mo*LOX, (**Figure 7A**), validate the MD structure as a reliable ground-state 3-D model of the active site ES complex.

### Comparison of the native MoLOX ES structure to that of Mn-SLO

The DAD between C11 of LA and the hydrogen acceptor on the Mn-bound oxygen of 3.4 ± 0.3 Å is the same as that determined from the X-ray crystal co-structure of coral 8*R*-LOX with arachidonic acid (3.4 Å for ‘C↔O’).^*8*^ Based on the comparison of the distribution of DADs, the ensemble of ground-state distances for LA in *Mo*LOX, as determined by a combination of ENDOR distance restraints and multiple MD trajectories, is ∼0.3 Å longer than in Mn-SLO, of 3.1 ± 0.2 Å (**Figure 9 and S9**).

**Figure 9.**
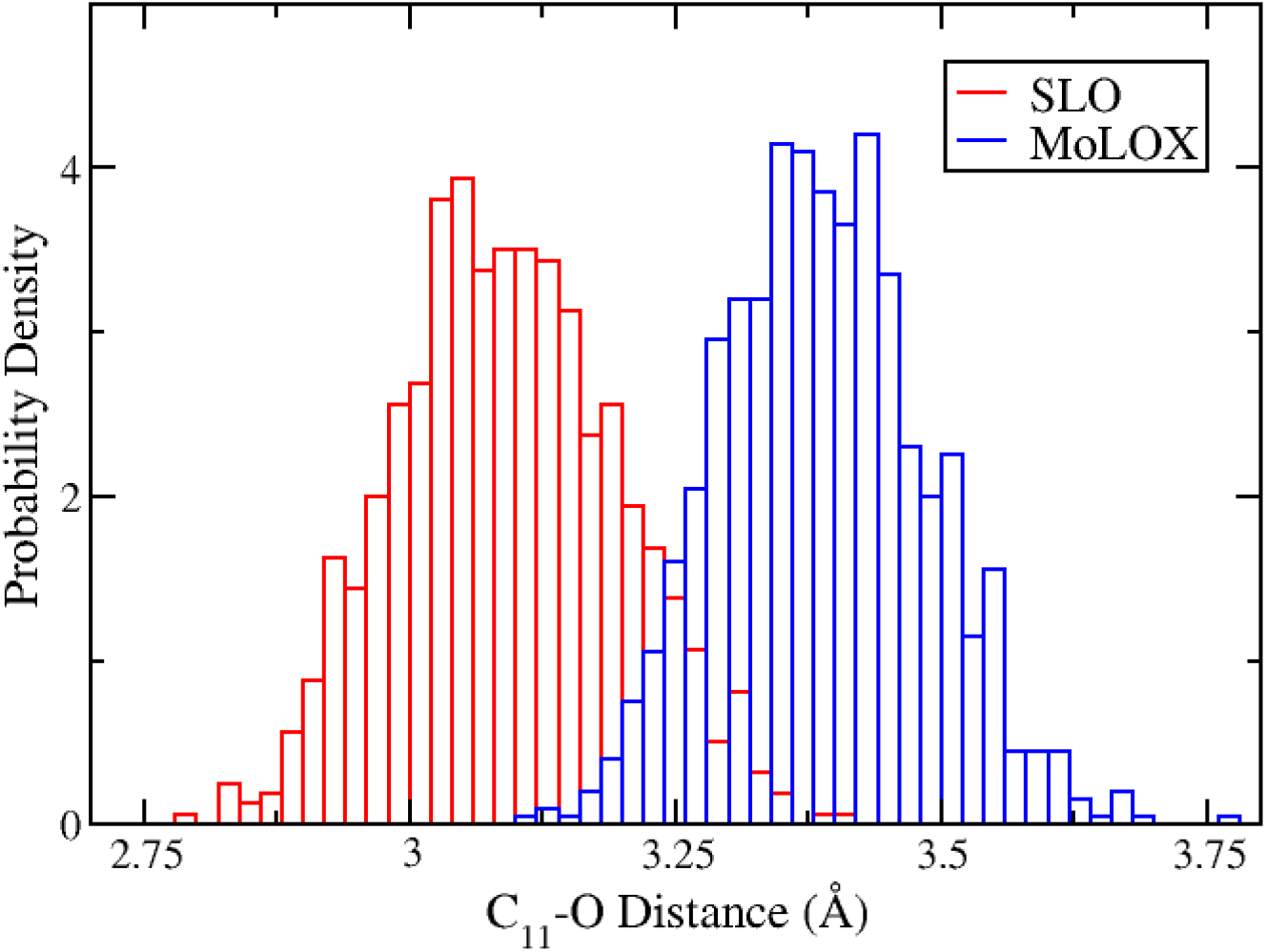
Ground-state DA distance distributions from ENDOR-guided MD trajectories for *Mo*LOX and SLO.

In the ENDOR-guided MD model of *Mo*LOX (**Figure 7A**), the carboxylate group of LA is positioned near the surface at the substrate entrance in a “carboxylate out” orientation and with hydrogen bonding interactions to two arginine residues, R525 and R528. Mutation of the Arg525 to alanine largely abolished catalytic activity for the reaction with the natural substrates (i.e. LA and ALA).^*16*^ While R528 mutational studies have not been reported, it is conserved across predicted fungal LOXs. Thus, R525 (and perhaps R528) of *Mo*LOX appears to help tether the carboxylate group of C18 fatty acid substrates, supporting a “carboxylate out” substrate binding orientation.^*16*^ In contrast, the binding orientation for LA in SLO is not completely resolved. Structural and kinetic studies using various substrates and analogues have provided evidence for both “carboxylate out” and “carboxylate in” models.^*37-39*^ One line of support for the “carboxylate in” model is the presence of an arginine residue, R707, buried at the end of the predominantly hydrophobic substrate channel of SLO. Mutation of Arg707 of SLO to leucine had a 3-fold reduction in the catalytic proficiency for the reaction with LA and a significantly lowered binding affinity for the product of LA, 13*S*-hydroperoxy-9,11-Z,E-octadecadienoic acid (13*S*-HPOD), which activates the enzyme.^*37*^ When modeling LA in the SLO active site using the “carboxylate in” model for MD simulations (**Figure S10**), the carboxylate group is within hydrogen bonding distance to R707.^*21*^ MD simulations on SLO were also previously conducted with LA in the “carboxylate out” model,^*21*^ but resulted in an unrealistic binding orientation with a significantly longer DAD of 4.1 Å and with C13 of substrate, which is the site for oxygen insertion,^*40*^ positioned away from the identified O_2_ channel. Thus, the “carboxylate in” orientation for LA in the SLO active site enables it to access catalytically relevant configurations.

These considerations indicate that LA adopts a ‘carboxylate in’ mode in SLO, but a ‘carboxylate out’ mode in *Mo*LOX (**Figure S10**). The difference in substrate binding is attributed to different orientation and structural alterations of the substrate channel shape,^*16, 41*^ which is influenced by sidechain bulk from primarily hydrophobic residues whose mutant kinetic properties are highlighted in **Tables 4** and **S1**. This difference in turn would strongly impact efforts to design inhibitors of *Mo*LOX.

### Relevance of ground-state DADs to the efficiency of hydrogen tunneling

Based on the multidimensional analytical rate model for enzymatic H-tunneling developed through decades of experimental and theoretical investigations centered on SLO, effective hydrogen wave function overlap in native H-tunneling enzymes is achieved in the so-called tunneling-ready state at compacted DADs of 2.7-2.8 Å.^*11, 42-45*^ In these reactions, the observation of a negligible differential activation energy between protium and deuterium transfer; Δ*E*_a_ = *E*_a_(D)-*E*_a_(H)) ≈ 0 for a native enzyme is a kinetic indicator of the effective achievement of tunneling-ready DADs without substantial additional contributions from local, isotope-dependent sampling motions involving the donor and acceptor atoms.^*22, 28*^ Both WT SLO and *Mo*LOX exhibit large ^D^*k*_cat_ and near-zero Δ*E*_a_ values for their rate-limiting hydrogen transfer reactions, which thus suggests compacted, optimized DADs at the tunneling-ready state. This compaction from the ground-state DADs, characterized by van der Waals distances of 3.1-3.4 Å, to TRS distances effective for tunneling, occurs transiently through multiple tiers of C11↔O distances, and facilitated by protein thermal motions involving a combination of the stochastic sampling along the conformational energy landscape and the environmental reorganization barrier (λ) along the reaction coordinate, which fine-tunes local geometries and electrostatics.^*11*^

Because the quantum-mechanical wavefunction overlap for hydrogen tunneling at the tunneling-ready state is inherently temperature-independent, the measured non-zero *E*_a_ in native enzymes reflects the contributions of thermal motions of the ES-complex that reduce the ground-state DAD to that in the tunneling-ready configuration. Thus, the observation of a longer ground-state DAD in *Mo*LOX compared to SLO, can explain why *E*_a_ for *Mo*LOX is larger than that of SLO: the longer DAD requires more extensive thermal protein motions to attain the optimized tunneling configuration, in which H-atom wavefunction density is equal on both D and A. This difference leads to *E*_a_ ≈ 10 kcal/mol for the reaction of WT *Mo*LOX with LA,^*14*^ whereas an anomalously low *E*_a_ ≈ 2 kcal/mol is observed for the same reaction in SLO.^*33*^ The link between the magnitude of the ground-state DADs and the energy needed for conformational activation (*E*_a_) is strengthened by the observed difference in the MD-derived root-mean square fluctuations (RMSF) given by the MD simulations for the positions of the substrate carbon atoms within the active sites of SLO and *Mo*LOX (**Figure S11**). SLO substrate carbons were found to exhibit RMSF of ∼ 1.5 Å for C1-C15, while the substrate LA undergoes larger fluctuations in *Mo*LOX during the MD trajectories, RMSF > 2.1 Å (**Figure S11**).

The MD trajectories for WT Mn-SLO,^*21*^ show that through stochastic conformational sampling the LA accesses C11-O distances as short as 2.8-2.9 Å (**Figure 9**, red), which nearly approach, but do not quite reach, DADs optimal for catalysis. Additional environmental reorganization (λ), then fine-tunes the DA distance and orientation to promote effective hydrogen wavefunction overlap for facile H-atom tunneling. Conversely, the shortest equilibrium DADs accessible through the analogous MD simulations of the *Mo*LOX-LA complex is only ∼3.1 Å (**Figure 9**, blue). This suggests that the *Mo*LOX ES complex requires additional isotope-independent sampling of the conformational energy landscape to achieve a rare, high-energy configuration(s) with shorter equilibrium DADs. This in turn implies that the probability of achieving a catalytically productive ES complex is diminished, leading to an expected decrease in catalytic rate (i.e., reduced *k*_cat_). This analysis is consistent with the trends in the experimental *k*_cat_ values for the *Mo*LOX and SLO reactions, with the former rate constant of 2 s^-1^ but a 100-fold greater rate constant for the SLO reaction. Alternatively, tunneling-ready C↔O distances for *Mo*LOX may also be achieved through a greater contribution from the environmental reorganization (λ_*Mo*LOX_> λ_SLO_).

This study elicits a fundamental question as to the extent that protein thermal motions can overcome unfavorable substrate positioning in the ground-state (not ideal distances for H-atom tunneling) in the case of evolutionarily related enzymes from different kingdoms, and how the magnitude of the activation energies for rate-limiting hydrogen transfer in enzymes also reflect these features. Future structural, kinetic, and theoretical studies on LOX orthologues will help quantify the contributions of the multi-dimensional coordinates to the activation energy barriers for hydrogen transfer across the lipoxygenase family to further understand how these enzymes have functionally diverged.

## Conclusions

The pathogenic fungal lipoxygenase, *Mo*LOX, contains a catalytically active, mononuclear Mn^2+^ metallocentre with coordination-sphere geometry distinct from that of the canonical iron LOXs, and that provides a natural electron-spin probe that is used here to examine the geometry of substrate binding by EPR/ENDOR spectroscopy. The combination with MD simulations yields a high-resolution model of the ground-state active-site structure of *Mo*LOX complexed with its native substrate, LA. The distance between the C11, which bears the hydrogen donor of LA, and the metal-bound oxygen (DAD) is ∼3.4 Å, in excellent agreement with an X-ray derived co-structure of 8*R*-LOX with AA. The ‘carboxylate-out’ binding mode of substrates bound to *Mo*LOX and 8R-LOX contrasts with the ‘carboxylate-in’ binding in the model plant lipoxygenase, SLO. Despite this difference, the *Mo*LOX DAD ∼3.4 Å is only slightly longer than the corresponding value for SLO (3.1 Å), which exhibits an anomalously low activation energy associated with hydrogen transfer. The structural and kinetic results presented herein also establish a relationship between the dynamical achievement of a geometry optimized for H-atom transfer in native LOX enzymes, and the degree to which subsequent protein thermal fluctuations are required to reach configurations favorable for catalysis because they exhibit enhanced hydrogen wave function overlap. The current work demonstrates the versatility of the ENDOR-guided MD approach to characterize the ground-state configurations of the substrate in the active site of the diverse and biologically important family of LOXs, which often resist X-ray crystallographic efforts. The structural information gleaned from this study, not least determination of the ‘carboxylate-out’ LA binding, is expected to guide future *in silico* screening efforts and rational inhibitor design of *Mo*LOX, an emergent target for taming the devastating rice blast disease.

## Materials and Methods

### Materials

Dideuterated and ^13^C labeled linoleic acid substrates were synthesized previously.^*48*^ All yeast/bacterial cells, media, salts, and buffers were purchased from Fisher Scientific, Sigma-Aldrich, or VWR at the highest grade possible.

### MoLOX Expression and Purification

WT *Mo*LOX was expressed in and purified from *P. pastoris* X-33 cells as previously described.^*14*^ The final purification for all variants was carried out using a HiPrep 26/60 Sephacryl S-200 column on an AKTA FPLC system with 50 mM HEPES (pH 7.5), 150 mM NaCl, and 10% glycerol. The fractions that corresponded to the peak with lipoxygenase activity were concentrated to 100 μM, frozen in aliquots with N_2_ (l), and stored at - 80°C. Mn content was determined using ICP-MS. Site-directed mutants were generated with the Qiagen QuickChange kit.

### EndoH Deglycosylation

An endoglycosidase H (EndoH) gene from *Streptomyces plicatus* was synthesized (GenScript) for recombinant expression in *E. coli* and subcloned in-line with his-tagged MBP construct (2CT-10 vector, Addgene plasmid #55209). Recombinant MBP-EndoH fusion was expressed in and purified from *E. coli* BL21 (DE3) cells. In brief, EndoH expression was induced by 1 mM IPTG once OD_600_ reached 0.6. Upon induction, the cultures were dropped to 18 °C and incubated overnight. Cells were lysed by sonication in lysis buffer (50 mM sodium phosphate, 100 mM NaCl, 8% glycerol, 2 mM MgSO_4_, pH 7.5, supplemented with lysozyme, DNAse I, and AEBSF). The protein was purified using NiNTA chromatography, dialyzed in 50 mM Tris, 150 mM NaCl, pH 8 buffer and stored at -80°C until use.

EndoH *Mo*LOX was prepared by treating WT *Mo*LOX with EndoH in a 20:1 (*Mo*LOX-EndoH) mass ratio. The reaction was carried out at 20°C overnight in 50 mM sodium acetate, pH 5.5. The optimal reaction conditions were determined using SDS-PAGE. In our hands, the reaction did not require α-mannosidase for complete digestion, as previously suggested.^*13*^ For large scale preparations, after the reaction, *Mo*LOX was separated from MBP-EndoH by passing the reaction over a NiNTA column equilibrated with 50 mM Tris (pH 8), 500 mM NaCl and 10 mM imidazole and then further purification using SEC FPLC. The removal of the glycans was confirmed by SDS-PAGE (**Figure S4**).

### Circular Dichroism (CD)

*Mo*LOX secondary structure and thermal stability was assessed by CD spectroscopy using a Jasco model J-815 CD spectrometer with bandwidths of 2 nm using a Starna cuvettes (path length of 0.1 cm). Samples were prepared at 3 μM in 25 mM potassium phosphate (pH 7). Under these conditions, the PMT voltage (HT) remained ≤ 600V in the range of 190-260 nm. Measurements for stability were also carried out with wavelengths set to 222 nm and the temperature range of 25-90°C (2°C intervals) with a temperature ramp up rate of 0.6°C/min.

### EPR/ENDOR Measurements

Q-band echo-detected pulsed EPR spectra (two-pulse echo sequence, π/2−τ−π−τ-echo) and Davies/Mims pulsed ENDOR spectra were collected on a spectrometer that has been described. All measurements were done with a helium immersion dewar, at 2 K. The Mims pulsed ENDOR sequence (three-pulse echo sequence, π/2−τ−π/2−T−π/2−τ−echo, with rf pulse (T_rf_) inserted in the interval, T, between second and third pulses) was used to probe the ^13^C hyperfine coupling of C10, C11 nuclei of labeled LA substrate. The Davies ENDOR sequence (three-pulse echo sequence, π/2-τ-π-T-π/2-τ-echo), with rf pulse (T_rf_) inserted in the interval, T, between 2^nd^ and 3^rd^ pulses) was used to measure ^1^H and ^14^N couplings of H_2_O and histidine coordinated to Mn^2+^. Simulation of EPR spectra was carried out using EasySpin. Simulation of ^13^C10, ^13^C11, ^1^H, ENDOR spectra were carried out using a purpose-written (*ad hoc*) program in labview.

### Orientation of Mn-C vector

The orientation of Mn-C vector in the ZFS coordinate frame is defined through Euler angles (θ, ϕ), where “θ” is the angle away from the unique ZFS axis (z) and “ϕ” defines the rotation in the x-y plane away from the ZFS x-axis. The orientation selection that allows determination of the orientation of **T** relative to the ZFS frame is provided by the ZFS-splitting of the EPR spectrum, which implies that for a labelled LA site, an ENDOR spectrum at each external field interrogates a defined set of orientations of the field relative to the Mn-^13^C vector, *r*. When the magnetic field is set to the low-field edge of the EPR spectrum, the signal arises only from the *m*_*x*_ -5/2 ↔ -3/2 manifold (**fig**. 1a) and one observes a single-crystal-like ENDOR spectrum with the field along the ‘*y*’ ZFS axis, while at the high-field edge one obtains an ENDOR spectrum from the same manifold, with the field pointing along the ‘*z*’ ZFS axis. ENDOR at each intermediate field is the sum of contributions from a well-defined set of orientations associated with the -5/2 ↔ -3/2 manifold, with the *m*_*s*_ *=* -3/2 ↔ -1/2 manifold contributing detectably.

### Docking and Molecular Dynamics

The initial structure of the WT *Mo*LOX enzyme was obtained from the PDB (code 5FNO^*16*^). The H++ webserver^*49, 50*^ was used to determine the protonation and to add hydrogen atoms under pH=9.0. Afterwards, the protonation states of the metal site residues were manually corrected if necessary due to the H++ webserver does not consider the metal ions or water molecules when determining the protonation states. The AutoDockVina program^*51*^ was used to dock the linolic acid (LA) into the protein binding pocket. A cubic box centered at the active site (with center has the coordinates [8.50 Å, 47.65 Å, 124.90 Å]) with length of 20 Å was used for the search space during the ligand docking. The *Mo*LOX model was represented by the AMBER12 forcefield^*52-55*^ with extensions for the LA ligand and Mn metal site. The partial charge for the LA^*56*^ and force constant of the Mn-ligand^*57*^ are taken from the literature. More details about the metal site parameters can be referred to our previous research.^*21*^

In the next step, hydrogen atoms were added and optimized the system. A periodic rectangular box of size 96 Å x 86 Å x 76 Å was used to solvate the protein-ligand system. The TIP3P water model was used to model the solvent molecules.^*58*^ 48 Na^+^ ions and 46 Cl^−^ ions were added to neutralize the system and provide a salt solution of NaCl with concentration of ∼0.13 M in the production runs. The following simulation procedure was performed for the protein-ligand system: (1) In the first equilibration step, with the protein-ligand system fixed, the solvent molecules and Na^+^ and Cl^−^ ions were minimized for 10000 steps and equilibrated for 100 ps at 300 K with a 2 fs timestep in the NVT ensemble. (2) Then the system was heated to 300 K in four steps 50K→100K→200K→300K. In each temperature steps, the system was minimized for 10000 steps and then propagated for 100 ps with a 2 fs timestep under 1 atm in the NPT ensemble. (3) Afterwards, two replicas were performed, for each has the system minimized for 20000 steps and then equilibrated for 20 ns at 300 K and 1 atm with a 2 fs timestep in the NPT ensemble, and finally has the production run for 20 ns at 300 K and 1 atm with a 2 fs timestep in the NPT ensemble. During Steps 2-3, the ENDOR restraints were applied to restrain the distance between Mn ion and C10/C11 as described in **Table 3**, with the force constant as 200 kcal/mol/Å^2^. Snapshots were saved every 40 ps during the production MD runs, yielding a total of 1000 frames for the final analyses. All the MD calculations were performed by using the NAMD-2.13b1 program.^*59*^

## Supporting Information

The Supporting Information is available free of charge on the ACS Publications website. Notes The authors declare no competing financial interest.

## Acknowledgments

The authors thank Prof. Mark Hoffmann (University of North Dakota) for assistance with MD data transfer and Prof. Anne Spuches (East Carolina University) for use of an anaerobic chamber for preparation of the EPR/ENDOR samples. This work used the Extreme Science and Engineering Discovery Environment (XSEDE), which is supported by National Science Foundation grant number ACI-1548562.^*60*^ Specifically, this work utilized the allocations granted to Tao Yu on Comet at the San Diego Supercomputer Center (SDSC) with the allocation number of CHE170073

