## Supporting info text for "^13^C ENDOR Spectroscopy-Guided MD Computations Reveals the Structure of the Enzyme-Substrate Complex of an Active, N-linked Glycosylated Lipoxygenase"

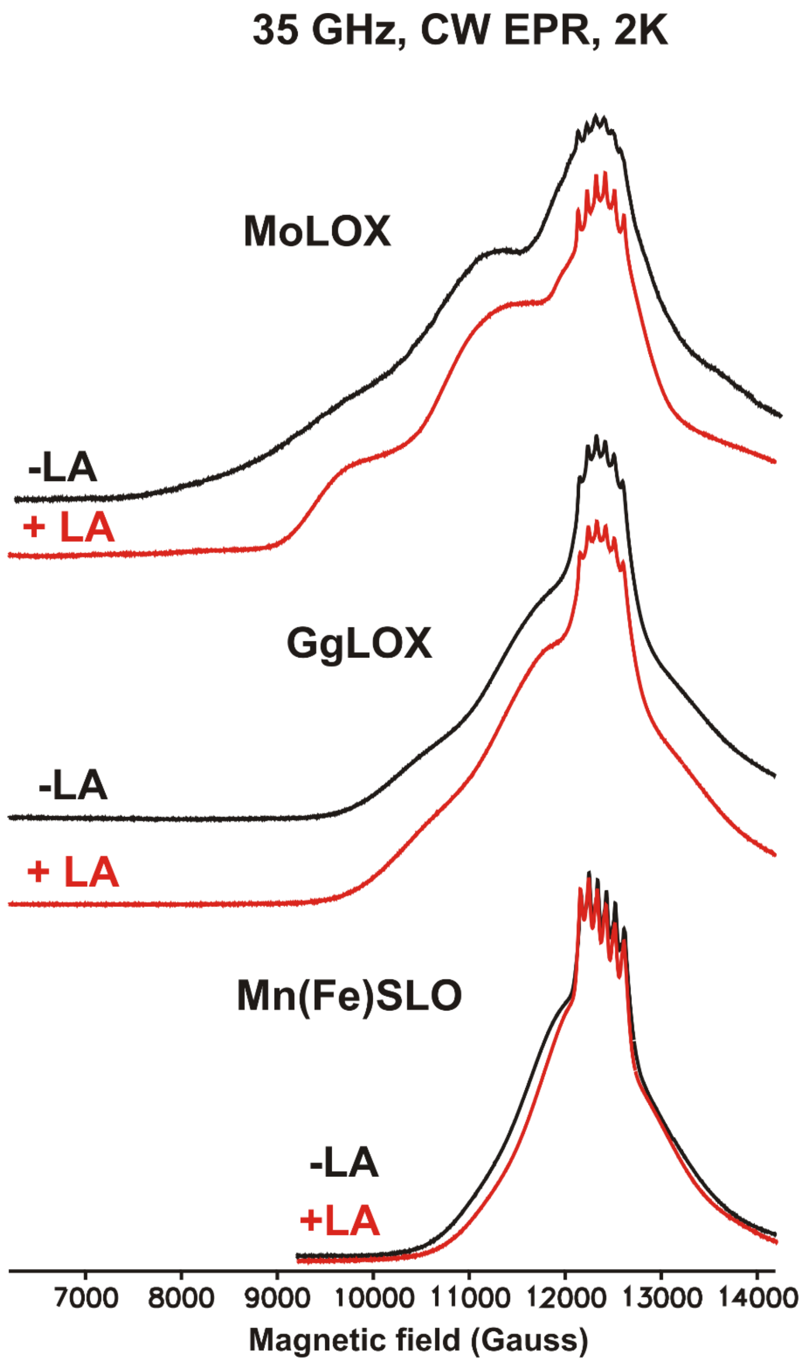


**Figure S1**: 35 GHz CW EPR spectra of wild-type *Mo*LOX, *Gg*LOX and Mn-SLO in the presence (in red) and absence (in black) of the substrate LA. Exp conditions: MW freq 34.9 GHz, MW power 1μW, Temp 2K

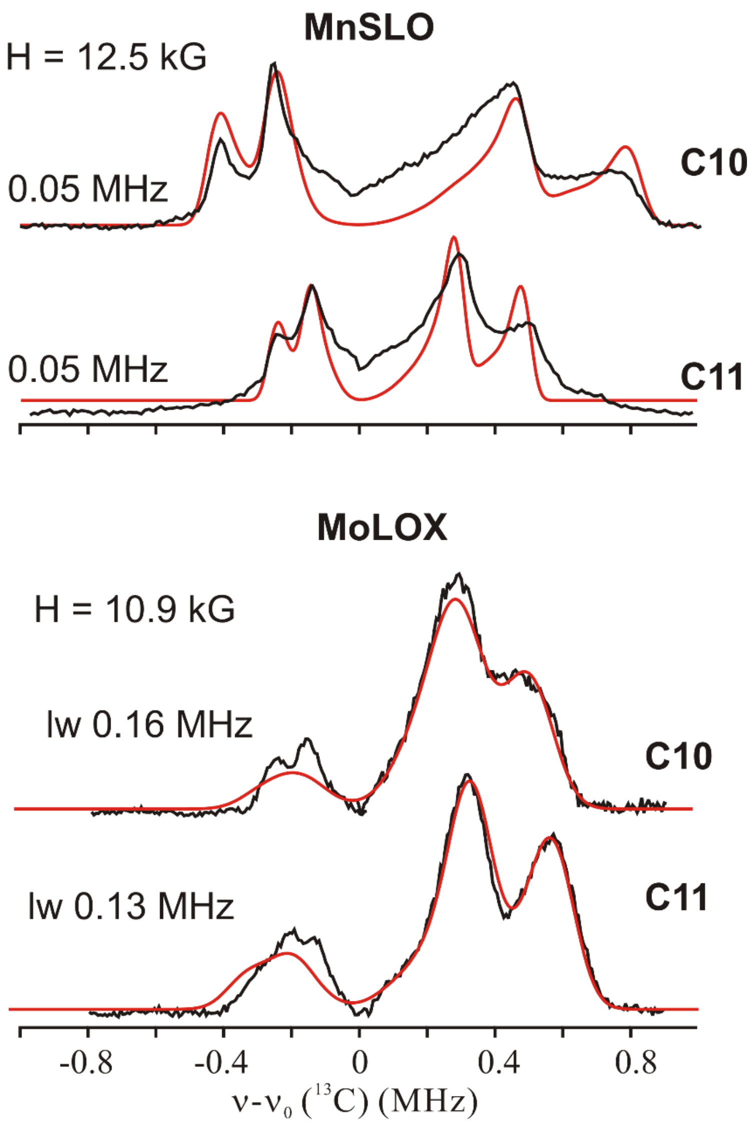


**Figure S2**: Comparing $^{13}C$ Mims ENDOR linewidths (lw) of C10, C11 of substrate LA between Mn-SLO and *Mo*LOX. Simulation in red. Exp conditions as in **Fig 3**.

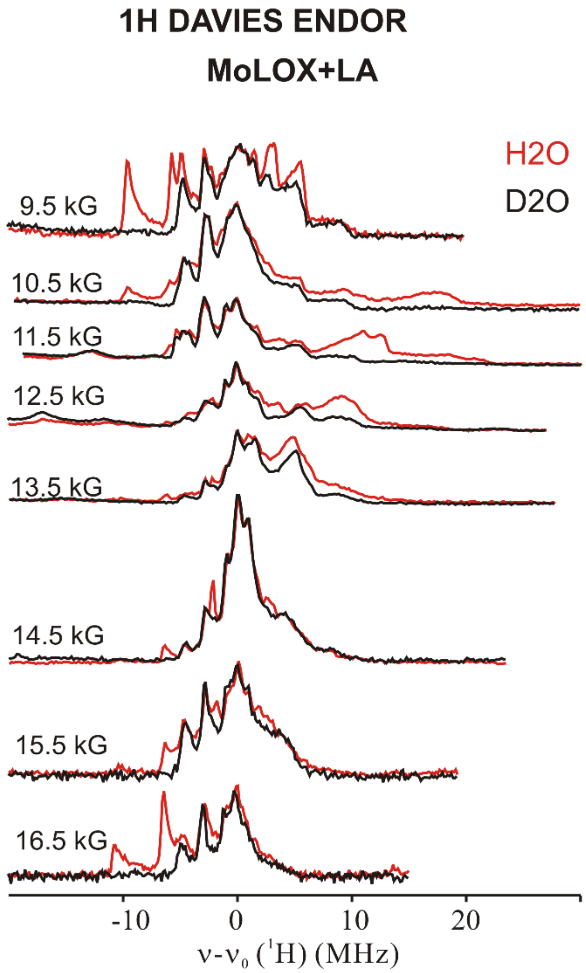


**Figure S3**: 35 GHz 2D field-frequency pattern $^{1}H$ Davies ENDOR for WT *Mo*LOX with LA in $H_2O$ buffer (in red) and in $D_2O$ buffer (in black). Conditions: microwave frequency ~ 34.8 GHz, MW pulse length ($\pi/2$) = 50 ns, $\tau$ = 500 ns, repetition rate =100 Hz, temperature 2 K.

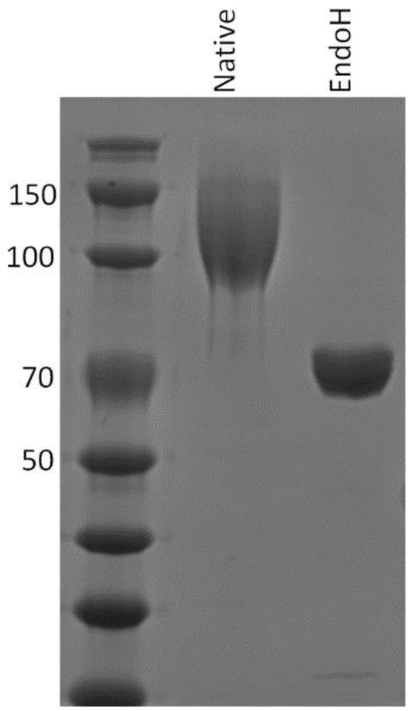


**Figure S4**. SDS-PAGE of wild-type and EndoH-treated *Mo*LOX. The molecular weights of the standards are listed along the left of the gel. The theoretical mass of *Mo*LOX based on primary sequence is 67 kDa.

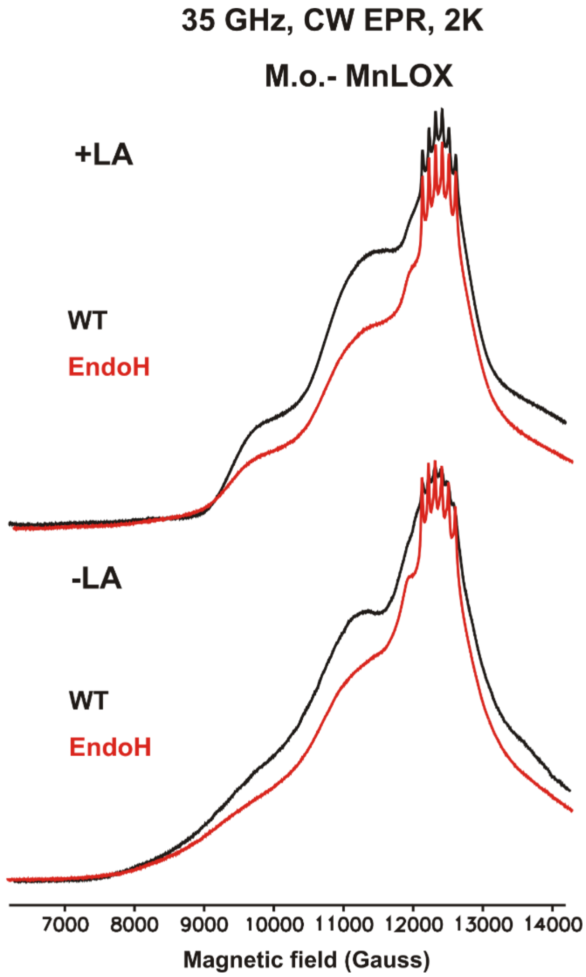


**Figure S5**: 35 GHz CW EPR spectra of *Mo*LOX, WT(black) and EndoH(red), in presence and absence of substrate LA. Exp conditions : MW freq 34.9 GHz, MW power 1μW, Temp 2K

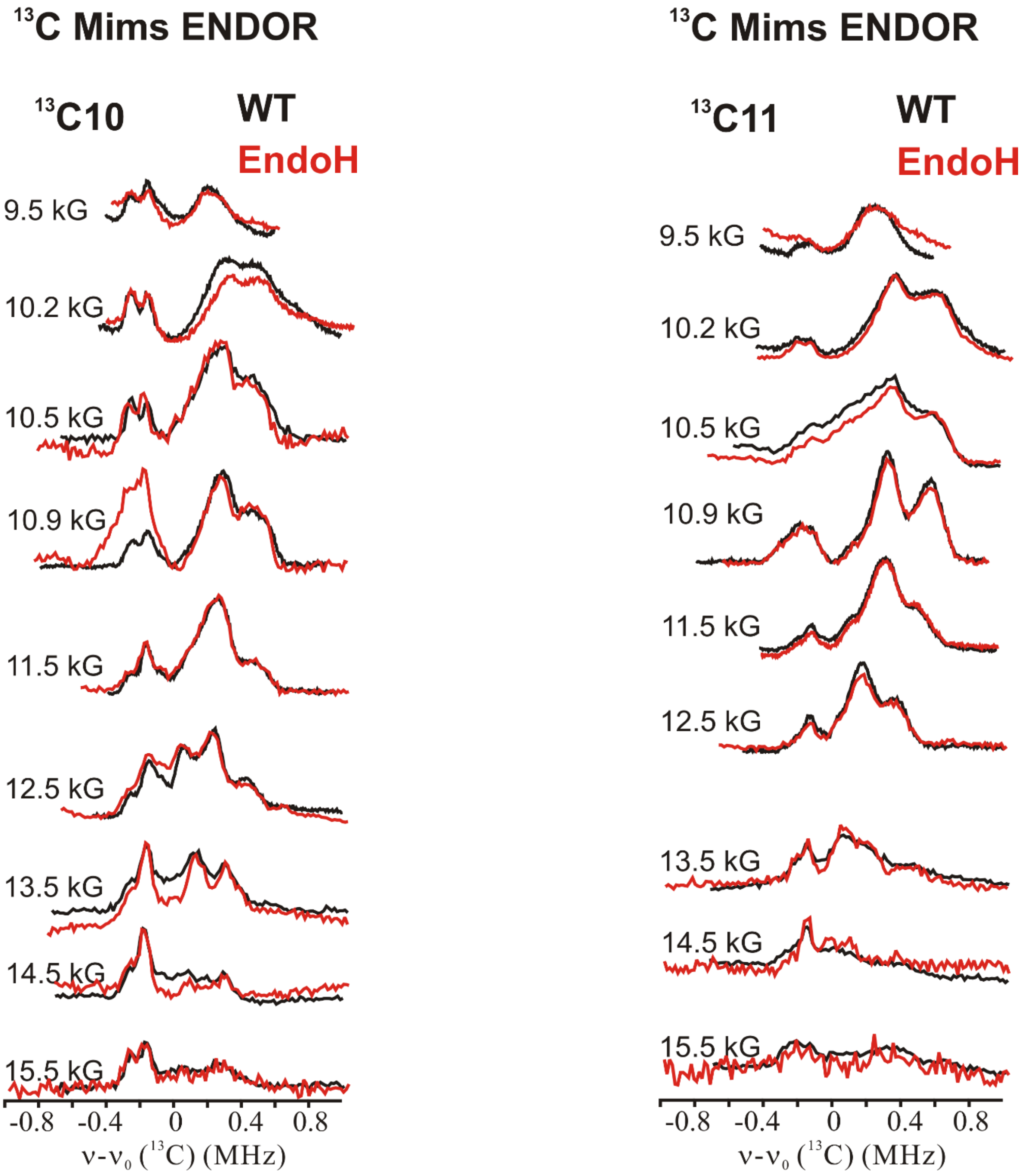


**Figure S6**: 35 GHz 2D field-frequency pattern $^{13}$C Mims ENDOR for *Mo*LOX WT (black) and EndoH (red) with $^{13}$C10-LA (left) and $^{13}$C11-LA (right). Conditions: microwave frequency ~ 34.8 GHz, MW pulse length ($\pi/2$) = 50 ns, $\tau$ = 1500 ns, repetition rate =100 Hz, temperature 2 K

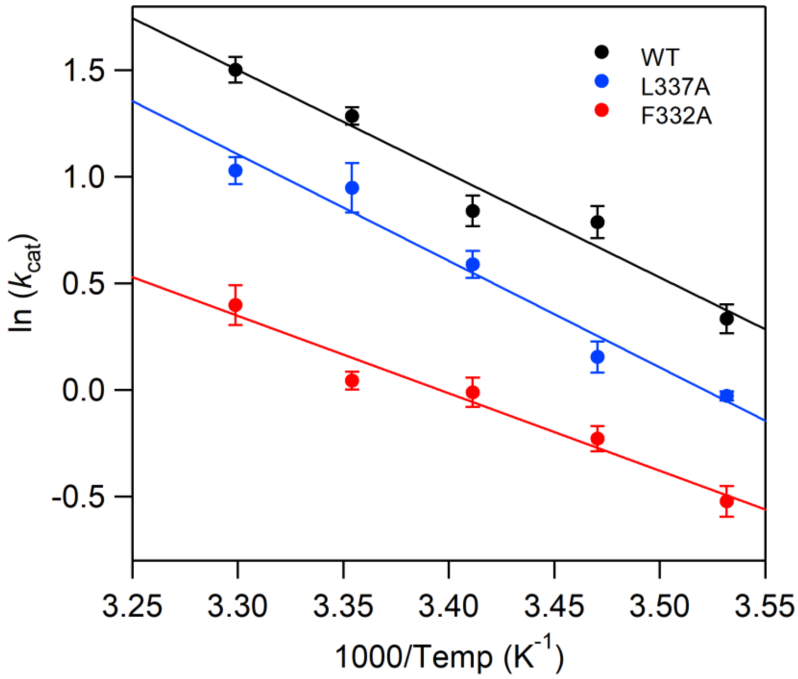


**Figure S7**. Arrhenius plots for WT (black), F332A (red), and L337A (blue) *Mo*LOX reactions with LA. The values represent the average of three independent experiments with the error bars representing s.e.m.

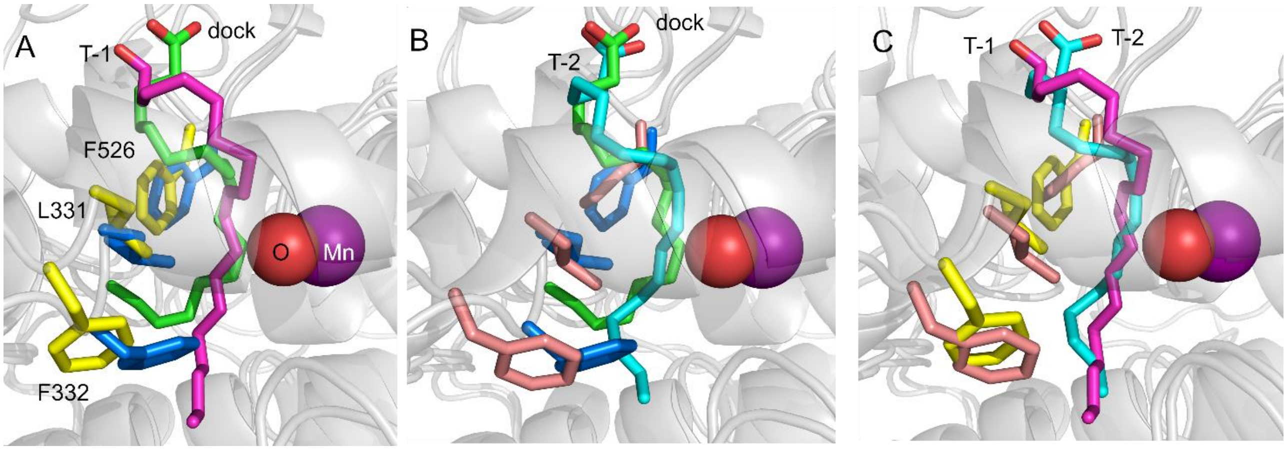


**Figure S8**. Structural overlays of the *in silico* docking model of LA (green) in *Mo*LOX (blue sidechains) (PDB: 5FNO) and the last snapshots of trajectory 1 (T-1; yellow sidechains, purple LA) and trajectory 2 (T-2; salmon sidechains, cyan LA) of the MD simulations.

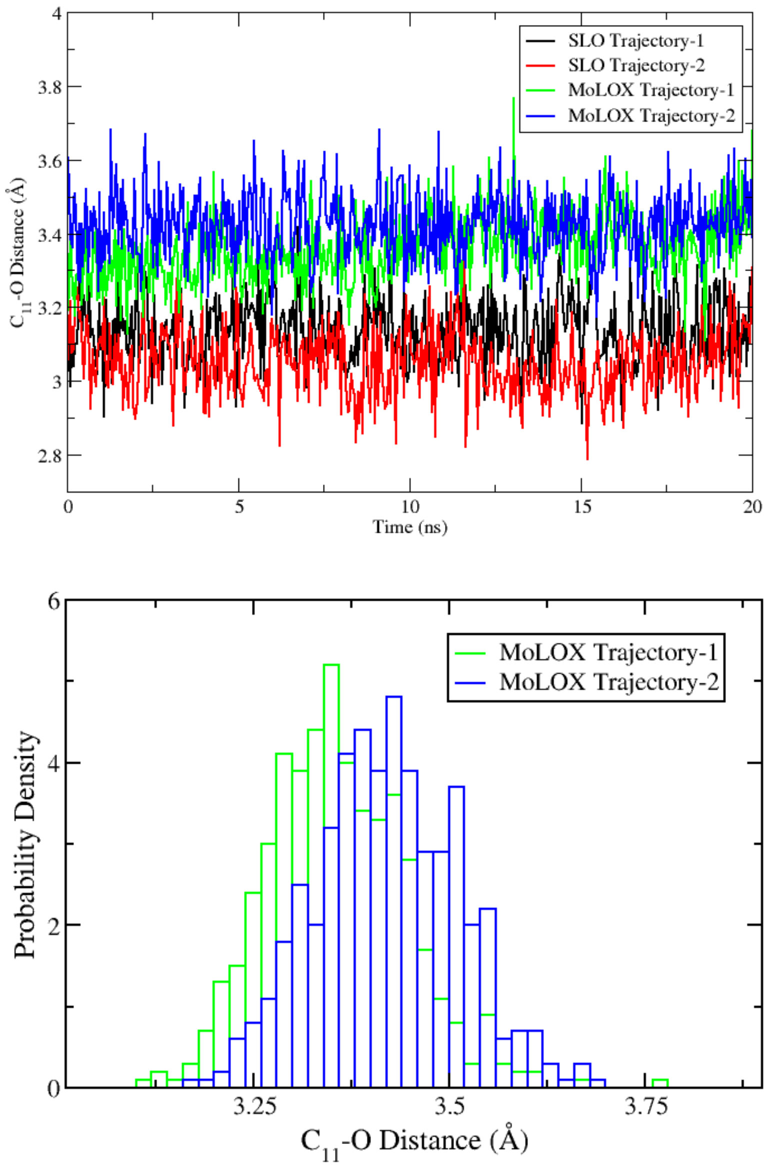


**Figure S9**. Calculated C11-O distance along initial two independent 20 ns molecular dynamic simulations of LA in Mn-SLO (red and black) and *Mo*LOX (green and blue).

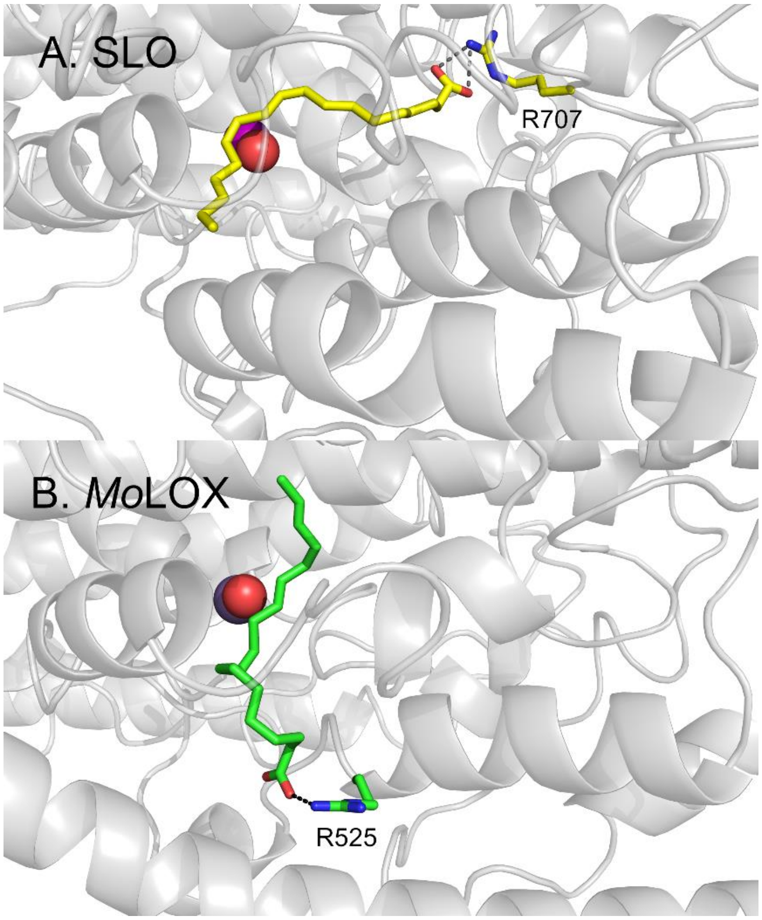


**Figure S10**. ES models of ‘carboxylate-in’ MnSLO-LA (A) and ‘carboxylate-out’ *Mo*LOX-LA (B), as determined by the ENDOR-guided MD approach.

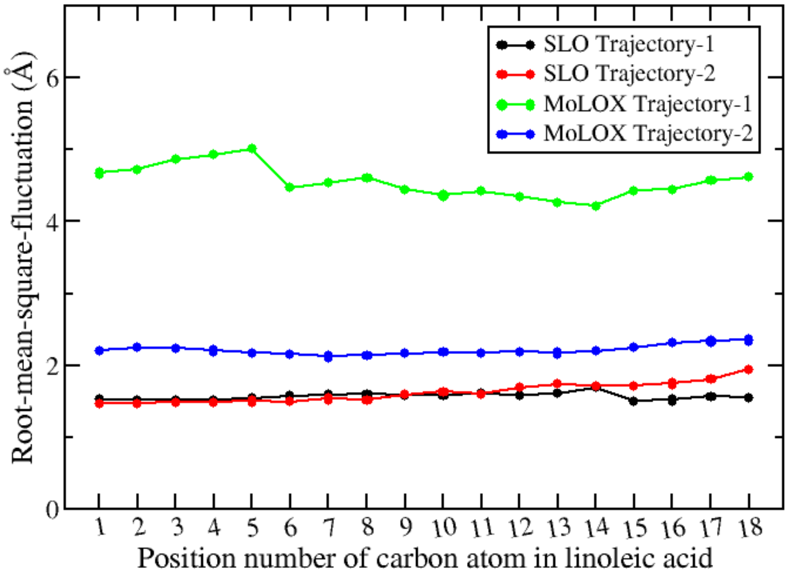


**Figure S11**. The root-mean-square-fluctuation (RMSF) values of the carbon atoms in the linoleic acid substrate for the MD trajectories of SLO and MoLOX. To calculate the root-mean-square-fluctuation (RMSF) values of the ligand carbon atoms along each trajectory, the following analysis was performed. First, the trajectory was aligned towards the first frame based on the protein backbone N, C$\alpha$, and C atoms. Afterwards, the average structure of the trajectory was obtained, with the trajectory was aligned towards this average structure based on the protein backbone N, C$\alpha$, and C atoms. Finally, the RMSF values of the ligand carbon atoms were calculated, with the RMSF of a specific atom *i* was calculated based on the following equation (herein *x* presents the atomic position): $RMSF_i = \sqrt{\langle(x_i - \langle x_i \rangle)^2\rangle}$. During the first trajectory of *Mo*LOX, the carboxylate oxygen atoms of LA forms more hydrogen bonds with the solvent water molecules and fewer hydrogen bonds with the Arg528 sidechain at the entrance of the substrate portal of *Mo*LOX (**Figure S12, Table S2**), which is linked to enhanced substrate motion in the *Mo*LOX active site as revealed by larger RMSF values for carbon atoms of the substrate during the first trajectory.

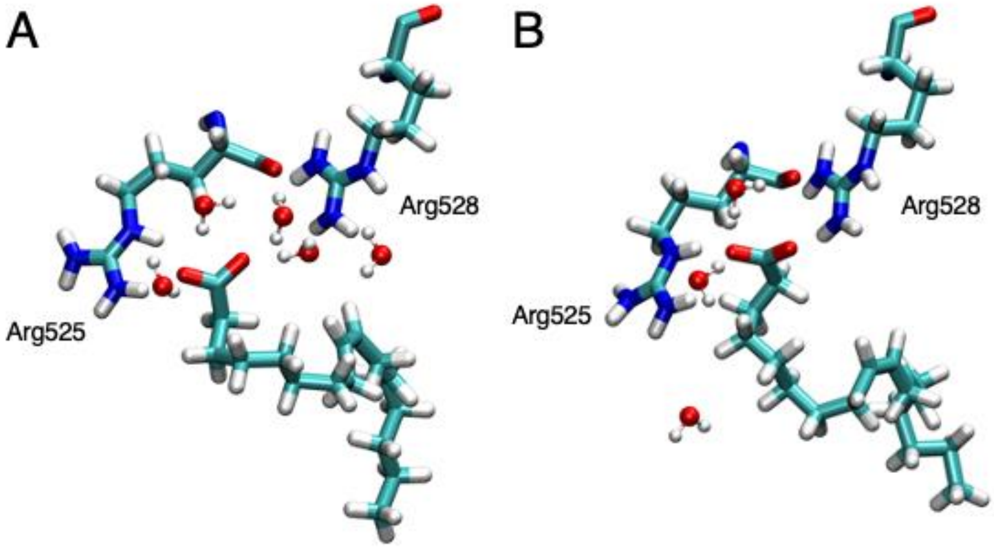


**Figure S12**. The last snapshot of the (A) first trajectory or (B) second trajectory. For clarity, only the linoleic acid, the two Arg residues at the entrance of the substrate binding site, and the water molecules within 3 Å of the linoleic acid are shown. Comparing to panel B, in panel A there are more water molecules and the water molecules disrupted the hydrogen bonding between the carboxylate group of linoleic acid and the Arg528 sidechain. Such a difference can also be seen in Table S2.

**Table S1. Comparative kinetics of *Mo*LOX mutants to SLO homologues.**

| *Mo*LOX variant | $k_{cat}$ ($s^{-1}$) | fold decrease | $E_a$ (kcal/mol) | SLO variant | $k_{cat}$ ($s^{-1}$) | Fold decrease | $E_a$ (kcal/mol) |
|---|---|---|---|---|---|---|---|
| WT | 2.1±0.2 | ---- | 9.5±0.5 | WT[a] | 297±12 | --- | 2.1±0.2 |
| L331A | 0.12±0.02 | 18 | N.D. | L546A[a] | 4.8±0.6 | 62 | 4.1±0.4 |
| F526A | 0.11±0.01 | 19 | N.D. | L754A[a] | 0.31±0.02 | 958 | 4.1±0.4 |
| F332A | 0.76±0.03 | 3 | 7.2±0.7 | L547[b] | N.D. | N.D. | N.D. |
| F338A[c] | N.D. | N.D. | N.D. | I553G[d] | 58±4 | 5 | 0.03±0.04 |
| L337A | 1.9±0.1 | unchanged | 9.9±1 | I552A[e] | 81±2 | 4 | -0.3±0.2 |
| L522A | 0.24±0.04 | 9 | N.D. | V750A[e] | 218±8 | 1.4 | 1.0±0.4 |

[a]From ref [1]

[b]No volume reducing sidechains constructed. Phe and Trp volume-increasing sidechain substitutions were constructed to test the impact of this residue on the predicted oxygen channel in SLO.[2]

[c]This mutant did not express.

[d]From ref [3]

[e]From ref [4]

**Table S2**. Fractions of the hydrogen bonds between the carboxylate group of linoleic acid and the Arg525 and Arg528 sidechains as well as the solvent waters along the two trajectories of the MoLOX systems. The following criteria were used to determine the hydrogen bond: donor-acceptor distance is within 3.0 Å and the donor-hydrogen-acceptor angle is within 135°.

| Hydrogen bond acceptor | Hydrogen bond donor | Trajectory 1 | Trajectory 2 |
|---|---|---|---|
| LIM@O2 | Arg525@NH2 | 46.8% | 80.8% |
| LIM@O1 | Arg525@NH2 | 43.6% | 0.0% |
| LIM@O1 | Arg525@NE | 32.6% | 0.0% |
| LIM@O2 | Arg525@NE | 27.6% | 66.6% |
| LIM@O2 | Arg528@NH2 | 7.6% | 0.0% |
| LIM@O1 | Arg528@NH2 | 3.4% | 99.4% |
| LIM@O1 | Arg528@NH1 | 0.0% | 27.4% |
| LIM@O2 | Arg528@NH1 | 0.4% | 0.0% |
| Sub Sum | | 162.0% | 274.2% |
| LIM@O2 | Water oxygen | 177.6% | 153.4% |
| LIM@O1 | Water oxygen | 157.2% | 0.0% |
| Sub Sum | | 334.8% | 153.4% |
| Total Sum | | 496.8% | 427.6% |
